# Combining genetic constraint with predictions of alternative splicing to prioritize deleterious splicing in rare disease studies

**DOI:** 10.1101/2022.02.28.482323

**Authors:** Michael J. Cormier, Brent S. Pedersen, Pinar Bayrak-Toydemir, Aaron R. Quinlan

## Abstract

**Background:** Despite numerous molecular and computational advances, roughly half of patients with a rare disease remain undiagnosed after exome or genome sequencing. A particularly challenging barrier to diagnosis is identifying variants that cause deleterious alternative splicing at intronic or exonic loci outside of canonical donor or acceptor splice sites.

**Results:** Several existing tools predict the likelihood that a genetic variant causes alternative splicing. We sought to extend such methods by developing a new metric that aids in discerning whether a genetic variant leads to *deleterious* alternative splicing. Our metric combines genetic variation in the Genome Aggregate Database with alternative splicing predictions from SpliceAI to compare observed and expected levels of splice-altering genetic variation. We infer genic regions with significantly less splice-altering variation than expected to be constrained. The resulting model of regional splicing constraint captures differential splicing constraint across gene and exon categories, and the most constrained genic regions are enriched for pathogenic splice-altering variants. Building from this model, we developed ConSpliceML. This ensemble machine learning approach combines regional splicing constraint with multiple per-nucleotide alternative splicing scores to guide the prediction of deleterious splicing variants in protein-coding genes. ConSpliceML more accurately distinguishes deleterious and benign splicing variants than state-of-the-art splicing prediction methods, especially in “cryptic” splicing regions beyond canonical donor or acceptor splice sites.

**Conclusion:** Integrating a model of genetic constraint with annotations from existing alternative splicing tools allows ConSpliceML to prioritize potentially deleterious splice-altering variants in studies of rare human diseases.

## Background

The diagnosis of Mendelian disease patients has progressed substantially owing to continued improvement of DNA sequencing technologies and software development for variant analysis. However, the diagnostic rate for these patients is still around 50%,^1–4^ leaving numerous patients with a diagnostic odyssey that persists indefinitely and results in financial, emotional, and physical burdens. Interpreting the effects of genetic variation outside of protein-coding exons is arguably one of the greatest challenges that remains. Discerning the impact of the non-canonical splice-site subset^5–9^ of these deleterious genetic variants is especially difficult. Consequently, potential splicing variants lying within introns or exons are commonly ignored during clinical diagnosis.

Up to 95% of all protein-coding genes in humans are predicted to undergo alternative splicing.^5,6^ Alternative splicing is a conserved process that expands the functional potential of nascent transcripts^10–12^ and requires tight regulation by the spliceosome, a large RNA and protein complex.^13–17^ Previous work has identified highly conserved DNA sequences that facilitate spliceosome assembly, function, and regulation.^10,16,18^ For example, variants at canonical di-nucleotide donor or acceptor splice sites lead to mis-splicing and are frequently deleterious. Disruption of alternative splicing is known to underlie numerous diseases.^8,9,19–24^ Therefore, variants impacting canonical donor or acceptor sites should be considered as strong evidence for pathogenicity according to the guidelines established by the American College of Medical Genetics (ACMG).^25^

Early experiments estimated that about 15% of disease-causing point mutations affect pre-mRNA splicing,^26^. More recent studies predict that between one-third to one-half of disease-causing variants disrupt splicing.^19,27–29^ These estimates are based on evidence of splice-altering variants occurring beyond the exon-intron junctions. Variants at important splicing motifs such as the intronic donor, acceptor, branch point, and polypyrimidine track sites would potentially lead to mis-splicing.^8,9,20^ Mis-splicing can also be caused by variants that affect splicing regulatory regions such as exonic and intronic splicing enhancers or silencers.^30–34^ Since most splicing occurs co-transcriptionally,^35–40^ variants that affect transcriptional processing may also alter splicing. For example, variants that alter the elongation rate and transcription kinetics of RNA polymerase II,^41–43^ RNA secondary structure,^44–47^, or nucleosome positioning^48,49^ could affect splicing during transcription. Furthermore, synonymous and non-synonymous variants in exonic regions affect splicing.^22^ Additional work has estimated that an individual exome-sequencing will harbor more than 500 potential splice-affecting variants of unknown significance50 and that up to 10% of the exonic disease-causing variants may also affect splicing.^22^ These findings suggest that current gaps in understanding, detecting, and interpreting splicing across the entire coding and non-coding body of protein-coding genes likely contribute to the low diagnostic rate of rare diseases.

Multiple software tools have been developed that improve the prediction and identification of splice-altering variants.^31,32,43,50–76^ For example, SpliceAI is a deep residual neural network that uses only the primary nucleotide sequence to yield accurate alternative splicing predictions.^76^ For each reference and possible alternative allele combination, SpliceAI provides four predictions. These predictions describe whether the variant may cause an acceptor loss, acceptor gain, donor loss, or donor gain. Similarly, SQUIRLS uses a Random Forest approach to predict whether a variant causes alternative splicing with similar performance as SpliceAI.^77^ SQUIRLS differs in that it uses various features such as conservation scores from phyloP78, sequence context parameters such as local distance from an exon and canonical splice sites, models of changes in spliceosome free energy binding, and other prediction tools such as ESRSeq.^79^ However, these and other existing approaches provide little to no guidance as to whether or not the variant is deleterious. Consequently, there is a need for improved methods that guide the prioritization of the subset of variants that may lead to deleterious alternative splicing to improve the diagnosis of rare diseases. This gap motivates our development of ConSpliceML, an ensemble machine learning method that combines regional measures of splicing constraint with computational predictions of the potential for genetic variants to cause alternative splicing. We demonstrate that ConSpliceML more efficiently prioritizes deleterious splicing variants in genic regions outside of canonical donor or acceptor sites.

## Results

### A statistical model of genetic constraint against deleterious splicing variants

We sought to develop a model of splicing constraint within protein-coding genes, motivated by the logic that owing to purifying selection, genomic regions leading to deleterious splicing will exhibit depletion of genetic variation in a large cohort of ostensibly healthy individuals. Using the single-nucleotide variants (SNVs) detected among 76,156 individuals in v3.1.1 of gnomAD^80^, we estimated substitution probabilities among protein-coding genes (**Methods**). We refined the substitution probabilities by estimating each substitution’s probability of leading to alternative splicing by binning the sum of the 12 SpliceAI scores provided for each reference nucleotide (**Supplementary Figure S1A, Figure 1A**). Each SpliceAI score estimates the potential for novel donor or acceptor sites if each of the three possible substitution alleles arose at a given reference position. Since SpliceAI’s deep neural network incorporates flanking sequence when modeling the cryptic splicing potential for each reference allele, its scores indirectly provide local sequence context to the resulting substitution probability matrix. As expected, we observe an overall decrease in the substitution probabilities as the sum of SpliceAI scores increases, indicating that variants predicted to alter splicing by SpliceAI are depleted in a healthy population (**Figure 1A, Supplemental Figure S1B**). However, since SpliceAI predicts whether a single allele will cause *alternative* splicing, our goal is to measure regional depletion of splicing variation to identify loci exhibiting genetic constraint against *deleterious* splicing.

**Figure 1:**
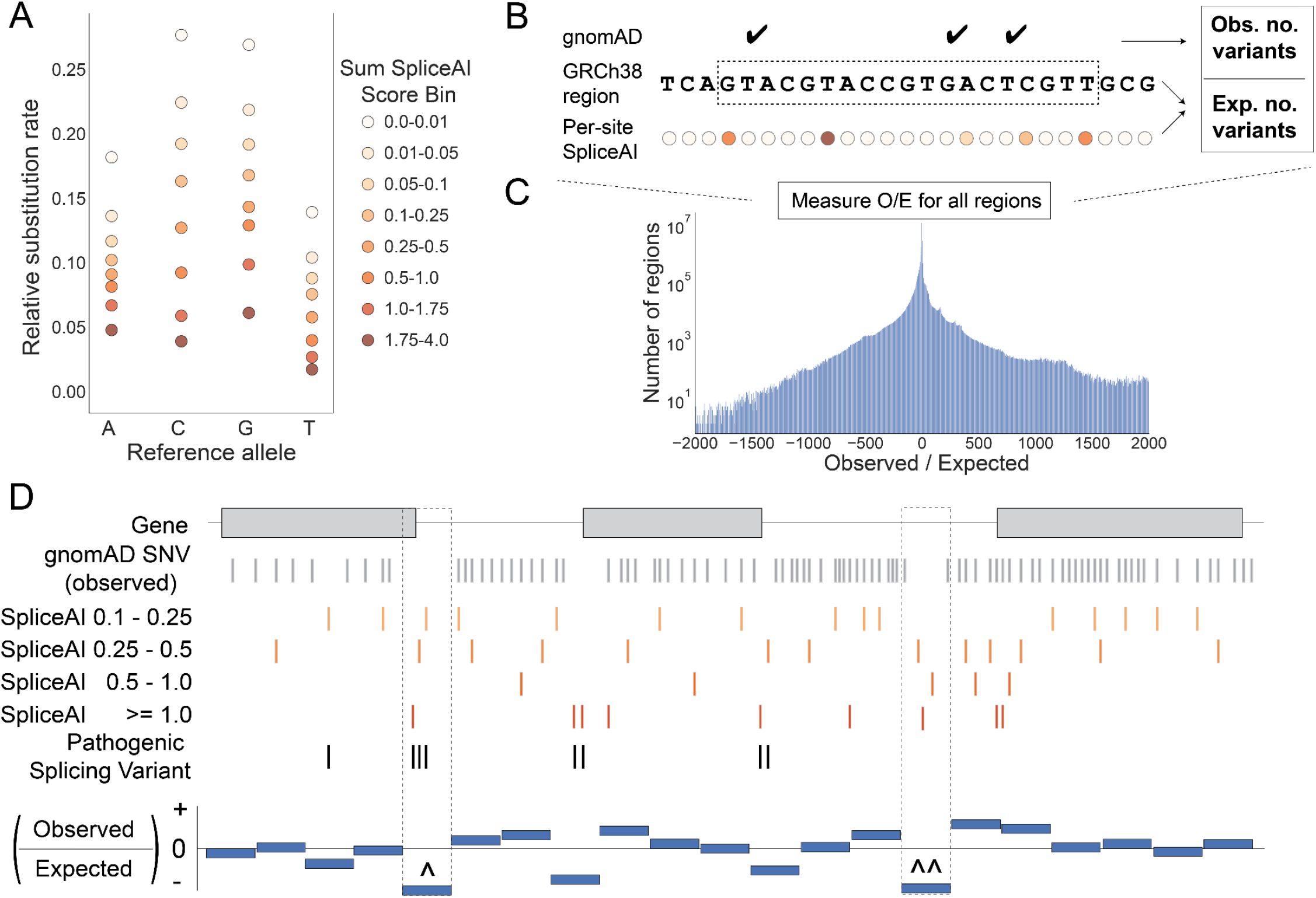
Overview of the regional splicing constraint model. (**A**) The per-site splicing substitution rate by reference allele and Sum SpliceAI score bin across autosomal protein-coding genes. The rate of no substitutions across all SpliceAI score bins for each reference allele is A>A = 0.9003, C>C = 0.8565, G>G = 0.8433, and T>T = 0.9347 (**B**) Calculating an Observed over Expected (O/E) ratio for a genomic region by counting the number of variants in that region from gnomAD and the number of expected variants with a given SpliceAI score. (**C**) The O/E score distribution. Smaller O/E scores indicate higher constraint against splicing, while larger O/E scores indicate lower constraints against splicing. (O/E plot truncated at −2000 to 2000 for visibility) (**D**) Representation of regional splicing constraint O/E scores across a hypothetical gene. The presence of gnomAD variants, in gray and the SpliceAI prediction for each position in the gene, in shades of red influences the splicing-specific observed and expected counts in a region. gnomAD variants with higher SpliceAI scores show evidence for more tolerance against splicing variation. In contrast, sites with a higher SpliceAI score and no gnomAD variant show evidence for less tolerance against splicing. Pathogenic splicing variants, in black, are commonly absent from gnomAD and have predictions of alternative splicing from SpliceAI. In this example, the regional constraint model identifies constraint signals at regions that harbor pathogenic splicing variants, such as at canonical splice regions (**^**) and cryptic splice regions (**^^**). All genomic positions in C without a SpliceAI score should be recognized as sites with a SpliceAI score < 0.1.

With this goal in mind, we compare the observed count of variation in gnomAD to what is expected (**Figure 1B**, **Methods**) across either entire genes or arbitrary regions within a gene. The expected counts for a given genomic region are derived from the substitution probability matrix (**Supplemental Table S1,2**), where the substitution probability for each nucleotide in that region is determined by the nucleotide’s reference allele, and SpliceAI summed score. The sum of the substitution probabilities for each nucleotide in the region yields the region’s expected substitution counts. We calculate a final ratio of the observed (O) to expected (E) variant counts that are further weighted based on each bin’s likelihood to contribute to alternative splicing events (**Figure 1C**, **Methods**). The null expectation for the O/E ratio was normalized to zero with an unbounded lower and upper limit to allow for a continuous spectrum of constraint (**Methods**). As a result, genomic regions less tolerant of splicing will have lower O/E ratios, as fewer splicing variants were observed in gnomAD than expected (a hypothetical example is depicted in **Figure 1D**). Finally, we transform O/E ratios to normalized scores ranging from 0.0 to 1.0, with 0.0 being the least constrained (tolerant) regions and 1.0 being the most constrained (intolerant) regions to splicing.

### Characterizing intolerance to aberrant splicing at the gene level

Gene-based genetic constraint scores such as the probability of loss-of-function intolerance (pLI)^81^ and LOEUF^80^ predict a gene’s intolerance to putative loss-of-function (pLoF) mutations. However, such metrics only include pLoF splicing mutations at the canonical donor and acceptor splice sites and are thus likely to miss deleterious splicing mutations. We were interested in characterizing each gene’s global intolerance to aberrant splicing mutations. Therefore, we measured splicing constraint using regions defined by the genomic interval defining each gene.

As a positive control, we tested expected patterns of constraint using gene sets known to underlie Autosomal Recessive (AR), Autosomal Dominant (AD), and Haploinsufficient (HI) disorders. Based on the evidence of strong or moderate sensitivity to heterozygous or homozygous pLoF variants,^80,82^ respectively, we hypothesized that the AD and HI genes would exhibit high predicted splicing constraint. In contrast, the distribution of splicing constraint for AR genes would be broader. We based these expectations on the notion that a single LoF variant arising on AD and HI genes may cause disease, while disease phenotypes on AR genes require LoF variants on both haplotypes. We included Olfactory Receptor (OR) genes as a negative control since these genes are known to be tolerant to pLoF variants.^83^ Since the majority of OR genes are single exons genes which are not expected to undergo splicing, we included genes shown to be non-essential for cell viability in CRISPR screens as a second negative control.^84^ As expected, both the AD and HI genes are more constrained against aberrant splicing; AR genes have a wide distribution across the score range, with OR genes and CRISPR non-essential genes less constrained (**Figure 2A**). Using several other gene sets, we identify similar patterns of expected splicing constraint in genes known to be involved in developmental delay and intellectual disability,^85–90^ genes shown to be essential in CRISPR screens,^84^ and genes shown to be tolerant to at least one homozygous pLoF mutation^81^ (**Supplemental Figure S2, S3**).

**Figure 2:**
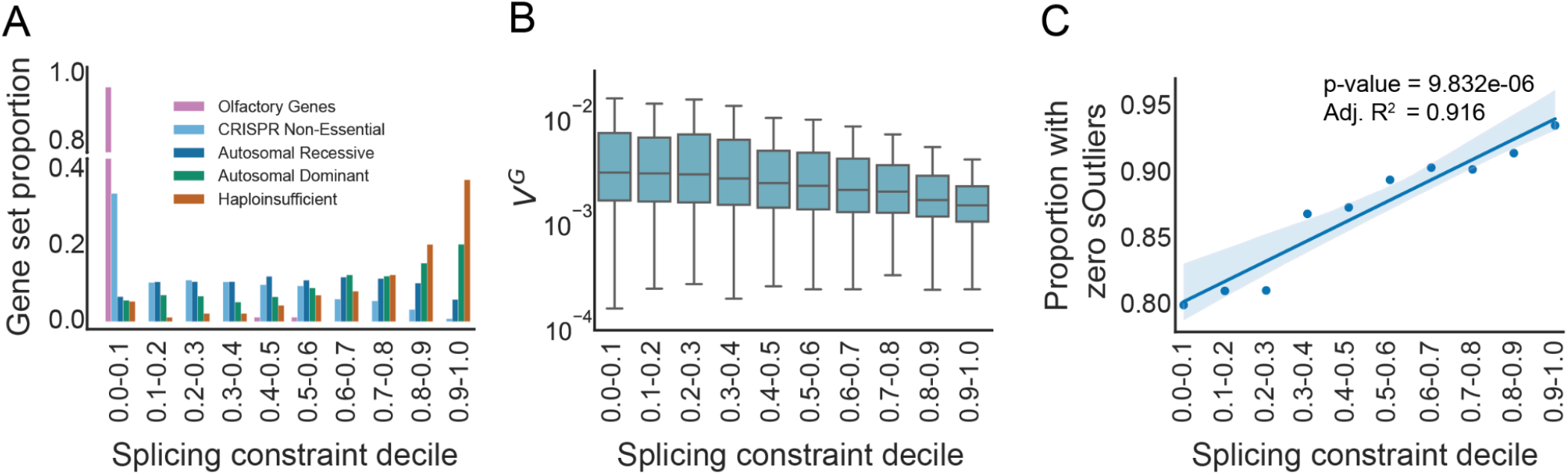
Patterns of genic splicing constraint. (**A**) The proportion of OR genes, CRISPR Non-essential genes, AR genes, AD genes, and HI genes across the splicing constraint deciles. Color corresponds to the gene set. (**B**) The distribution of V^G^ values by splicing constraint deciles (p-value = **2.488e-31)**. Significance determined using ordinary least squares linear regression. The boxes’ interquartile range (IQR) ranges from the 25th to 75th percentile. The horizontal black line in each box represents the median V^G^ value for that decile. The whiskers are 1.5X the IQR. Outliers are not plotted. (**C**) The proportion of genes in each splicing constraint decile that had zero significant sOutliers (p-value = **8.832e-06**, Adjusted R^2^= 0.916). Significance determined using ordinary least squares linear regression. The dark blue line represents the linear regression line. Light blue shade represents the 95% confidence interval. OR = Olfactory Receptor, AR = Autosomal Recessive, AD = Autosomal Dominant, HI = Haploinsufficient, sOutliers = splicing outliers.

The patterns of splicing constraint for these gene sets are strongly correlated with the constraint on coding sequence measured by LOEUF and moderately correlated with pLI (**Supplemental Figure S4**). We expect such correlations since genes under selective pressure should be broadly intolerant of any pLoF variant. For example, a pLoF nonsense variant in a gene with an AD inheritance pattern should have a similar constraint against a pLoF variant that leads to aberrant splicing. However, we also expected differences in constraint patterns since the splicing constraint model combines constraint in both coding and non-coding regions of a gene, whereas LOEUF and pLI focus solely on coding regions. Additionally, we anticipated the observed differences in constraint as the splicing constraint model is based on splicing predictions while pLI and LOEUF integrate all types of pLoF variation (**Supplemental Figure S4**).

Given that deleterious, splice-altering variants commonly trigger nonsense-mediated decay (NMD),^91^ which often reduces gene expression, we compared our gene-wide splicing constraint estimates to a measure of dosage sensitivity. We hypothesized that a gene’s sensitivity to changes in expression would be correlated with a gene’s constraint against splicing. To test this hypothesis, we used the expected genetic variation in gene expression (V^G^)^92^ calculated from the RNA-seq expression data in the Genotype-Tissue Expression (GTEx) project^93,94^ to measure a gene’s sensitivity to expression variation. V^G^ measures each gene’s expression variance and represents how tolerant a gene is to genetic variants that change expression. As expected, splicing constraint is significantly negatively correlated with V^G^ (p-value = 2.488e-31, **Figure 2B)** and remains significant after controlling for the number of exons per gene (p-value = 5.99e-26, **Supplemental Figure S5**). Genes in the highest constraint decile have a mean of 4.23-fold less variance in gene expression than genes in the lowest decile. Furthermore, the distribution of V^G^ is significantly different across splicing constraint bins after FDR correction (**Supplemental Figure S5**).

Since most genes are alternatively spliced, we hypothesized that genes less tolerant to variation in alternative splicing events would have a higher genic splicing constraint than genes more tolerant to alternative splicing variation. We used alternative splicing events detected in GTEx by Ferraro et al.^95^ to test our hypothesis of observing a depletion in alternative splicing for genes under higher splicing constraint. To estimate alternative splicing tolerance, we used the alternative splicing events that were significant outliers from the population distribution of alternative splicing events in a gene (sOutliers). As expected, there is a significant positive correlation with the proportion of genes having zero significant sOutliers and the splicing constraint for those genes. This observation indicates that genes more intolerant to alternative splicing have significantly fewer sOutliers (p-value = 8.83e-06, **Figure 2C, Supplemental Figure S6**). Additionally, we observe a negative correlation between splicing constraint and the proportion of genes having at least one significant sOutlier, indicating genes more intolerant to alternative splicing have significantly fewer sOutliers (p-value = 8.83e-06, **Supplemental Figure S6**).

### A regional model of splicing constraint

Genic loci outside of canonical splice junctions are involved in splicing.^8,12,18,23,39,50,96^ While a gene-wide measure of splicing constraint reveals global constraint against splicing, it is insufficient for identifying the focal origins of constraint, especially when motifs drive the signal outside of exon-intron junctions. For example, we expect the highest levels of splicing constraint to be localized to regions of genes that are critical to splicing. It is thus reasonable to consider an intragenic regional model of constraint that identifies focal regions within a gene that are most intolerant of aberrant splicing. Indeed, regional constraint metrics such as Constrained Coding Regions (CCR)^97^ highlight regional variability in constraint against pLoF variation within protein-coding genes. In the case of diagnosing rare diseases, it is imperative to identify and prioritize the deleterious causal variant, and using a regional level metric allows one to estimate where causal splicing variants would lie within a gene. Therefore, we created multiple regional splicing constraint models using various window sizes to determine the optimal region size to predict the pathogenicity of a splicing variant. We used window sizes that balanced smaller sizes with more spatial resolution with larger sizes that would contain more expected variants and therefore give more power to detect a difference between observed and expected splicing variation (**Methods**, **Supplemental Figure S7**).

As a positive control, we evaluated the regional splicing constraint model at various region sizes around exon features using a set of constitutive exons (exons present in every isoform of a gene) and cassette exons (exons alternatively spliced in or out of a gene depending on the expressed isoform) defined by Busch et al.^98^ We hypothesized that the splicing constraint at exons should generally be higher than in the surrounding 5’ and 3’ introns of those exons. This hypothesis comes from the fact that coding regions, in general, are more constrained than non-coding regions and that the exon boundaries are determined by splicing definition. As expected, we observe an increase in constraint as we move towards the 5’ end of an exon and a decrease in constraint as we move away from the 3’ end (**Figure 3A, Supplemental Figure S8**). Additionally, as expected, we see that a genetic constraint model unaware of splicing has a similar constraint pattern around exons. However, the change in constraint is much less dramatic than the splicing aware constraint model (**Supplemental Figure S9**). This observation further suggests that the model captures specific constraints beyond the difference in constraint between coding and non-coding regions of genes. Finally, we observe differences in the magnitude of constraint-driven by the inclusion rate of an exon, where exons with a higher inclusion rate are slightly more constrained than exons with a moderate rate. This observation supports previous high-inclusion cassette exons being characteristically identical to constitutive exons.^98,99^

**Figure 3:**
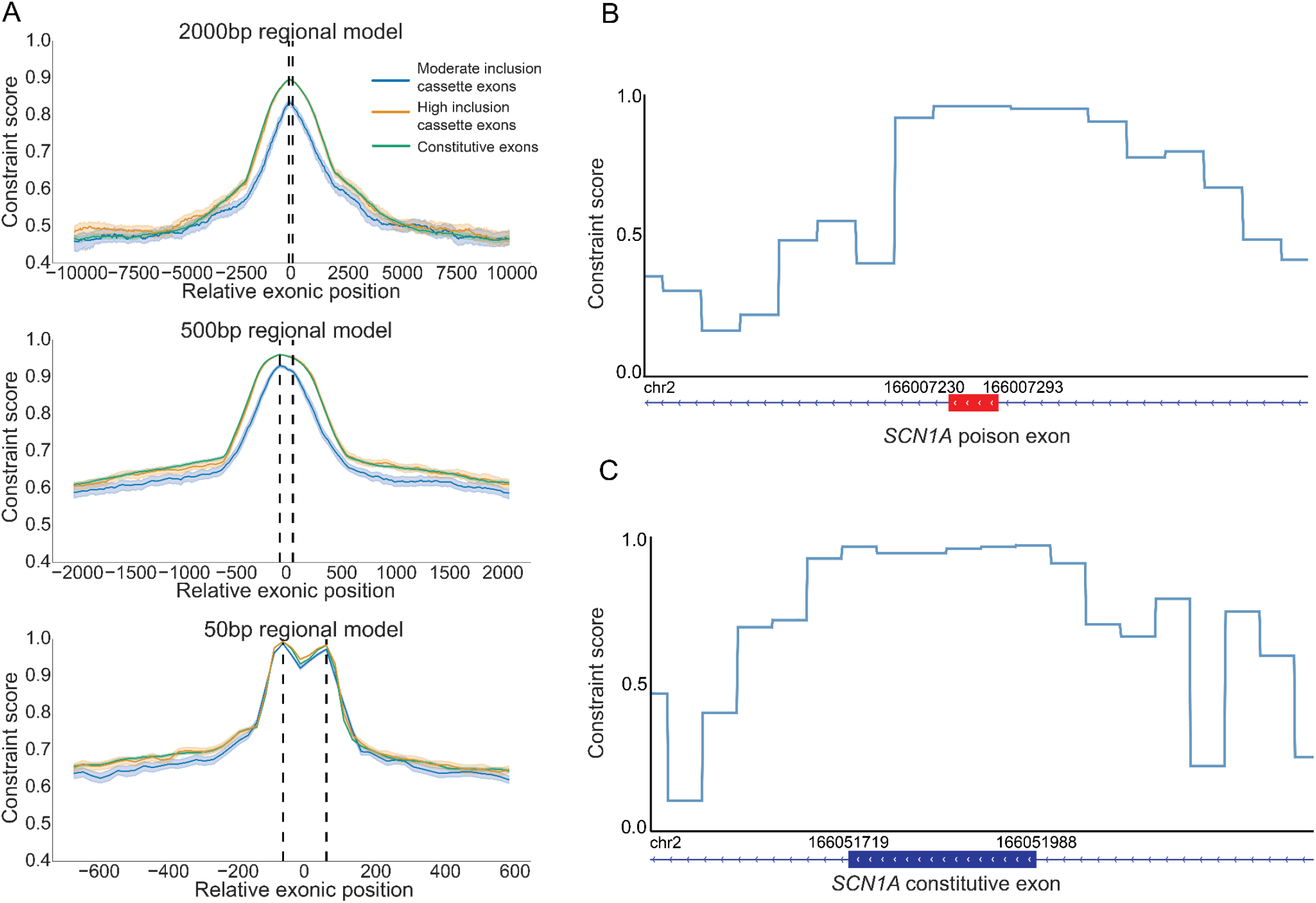
Splicing constraint around exon features. (**A**) The pattern of splicing constraint at and around exon features including constitutive exons (Full Inclusion Exons) and cassette exons (Moderate Inclusion Exons and High Inclusion Exons). Full Inclusion Exons have an inclusion rate of 100%. High Inclusion Exons have an inclusion rate >= 90% and < 100%. Moderate Inclusion Exons have an inclusion rate < 90%. Plots are oriented 5’ to 3’ from left to right. Vertical dotted lines represent the relative start (left) and end (right) position of exons. Colors represent exon class based on inclusion rate. Dark lines for each exon class represent the median regional constraint score along the relative region around exons. The lighter shaded regions around the line represent the 95% CI determined by 1000 bootstrap iterations for each exon class. Oriented by the window size with the 2000bp model on the top, 500bp model in the middle, and 50bp model on the bottom. Figure color legend is located to the right of the top plot. (**B,C**) Regional constraint score profiles for a poison exon in (B) and a randomly selected canonical *SCN1A* exon (C). The red horizontal bars in B represent the poison exon, and the dark blue horizontal bars in C represent the canonical exon. The gene track around the exon feature is shown below the regional constraint score profile with the GRCh38 3’ and 5’ (left and right, respectively) genomic coordinates for that exon feature above the gene track. *SCN1A* is on the negative strand; therefore 5’ to 3’ is from right to left in B and C. The regional constraint score track represents the splicing constraint profile using the 50bp regional model.

We note the difference in constraint profiles based on the size of the constraint region. As the region size decreases, the difference between exon classes as a whole diminishes, while the specificity of the feature driving the constraint increases. For example, we see an increase in the specificity of the 5’ and 3’ canonical splicing motifs as we move from the 2000bp splicing constraint model to the 50bp model (**Figure 3A**, **Supplemental Figure S8**). This result indicates that we can define an optimal region size based on the feature of interest. Splicing is primarily driven by small regulatory motifs such as the canonical acceptor and donor motifs, the polypyrimidine tract, and enhancer or silencer motifs. Therefore, an ideal region size would capture splicing motifs while excluding irrelevant adjacent nucleotides around the motifs that do not participate in splicing. However, the region also needs to encompass a sufficient number of nucleotides to calculate a robust O/E signal. Therefore, the region’s size needs to balance the size of a splicing motif and a region with a sufficient number of nucleotides to establish a robust O/E signal (**Supplemental Figure S7, Methods).** Empirically testing this hypothesis further supports a smaller region size as the optimal size for detecting focal splicing constraint within genes. Specifically, we found that the 50bp ConSplice model was sufficiently large to encompass splicing motifs and capture O/E signal, while also small enough to identify focal constraint, likely driven by the splicing motifs (**Figure 3A, Supplemental Figures S7,8,10**, **Methods**).

Beyond annotated exons, the regional splicing model also captures constraint against splicing at poison exons, which are conserved, alternatively spliced exons that lead to nonsense-mediated decay (NMD) caused by a premature termination codon located in the poison exon.^100^ While many poison exons are important regulatory features of the genome, others, when expressed, cause deleterious effects.^100^ For example, multiple deleterious poison exons have been found in *SCN1A,* leading to Dravet Syndrome (DRVT [MIM: 607208]), a severe neurodevelopmental disorder marked by epileptic encephalopathies.^96,101,102^ The 50bp regional model detects each of the three poison exons in *SCN1A* with constraint scores at or above 0.95. The poison exons rank in the top 8^th^ percentile and are in the top 2% of noncoding regions in *SCN1A.* Additionally, two of the three poison exons have higher constraint scores than the median constraint seen in coding regions of *SCN1A,* with the third having a constraint score of 0.017 below the median. (**Supplemental Figure S11**). These poison exons also have similar constraint profiles as the annotated exons in *SCN1A* (**Figure 3B,C; Supplemental Figure S12; Supplemental Table S3**), which replicates the pattern seen for constitutive and cassette exons (**Figure 3A**).

### Interpreting pathogenic splicing variation with a regional model of splicing constraint

We next evaluated how well the regional splicing constraint model was able to identify pathogenic splice-altering variants. We curated 376 pathogenic splice altering variants from literature, where each variant has some functional support for its effect on splicing (**Supplemental Table S4**). Additionally, each variant was associated with a disease, such as Hereditary Hemorrhagic Telangiectasia (HHT2 [MIM: 600376]),^103–110^ Stickler Syndrome(STL1 [MIM: 108300]),^111–115^ Optic Atrophy 1 (OPA1 [MIM: 165500]),^116,117^ Marfan Syndrome (MFS [MIM: 154700]),^118,119^ Retinoblastoma (RB1 [MIM: 180200]),^120,121^ and Neurofibromatosis type 1 (NF1 [MIM: 162200]).^122,123^ The regional constraint model placed 86% of the 376 variants at or above a constraint score of 0.95, indicating strong constraint against alternative splicing. For example, it classified 89% of intronic non-canonical splice-altering variants in *ACVRL1,* a gene that has well-characterized intronic splice-altering variants that cause Hereditary Hemorrhagic Telangiectasia (HHT2 [MIM: 600376]), with constraint scores at or above 0.88 (**Figure 4A-C**).

**Figure 4:**
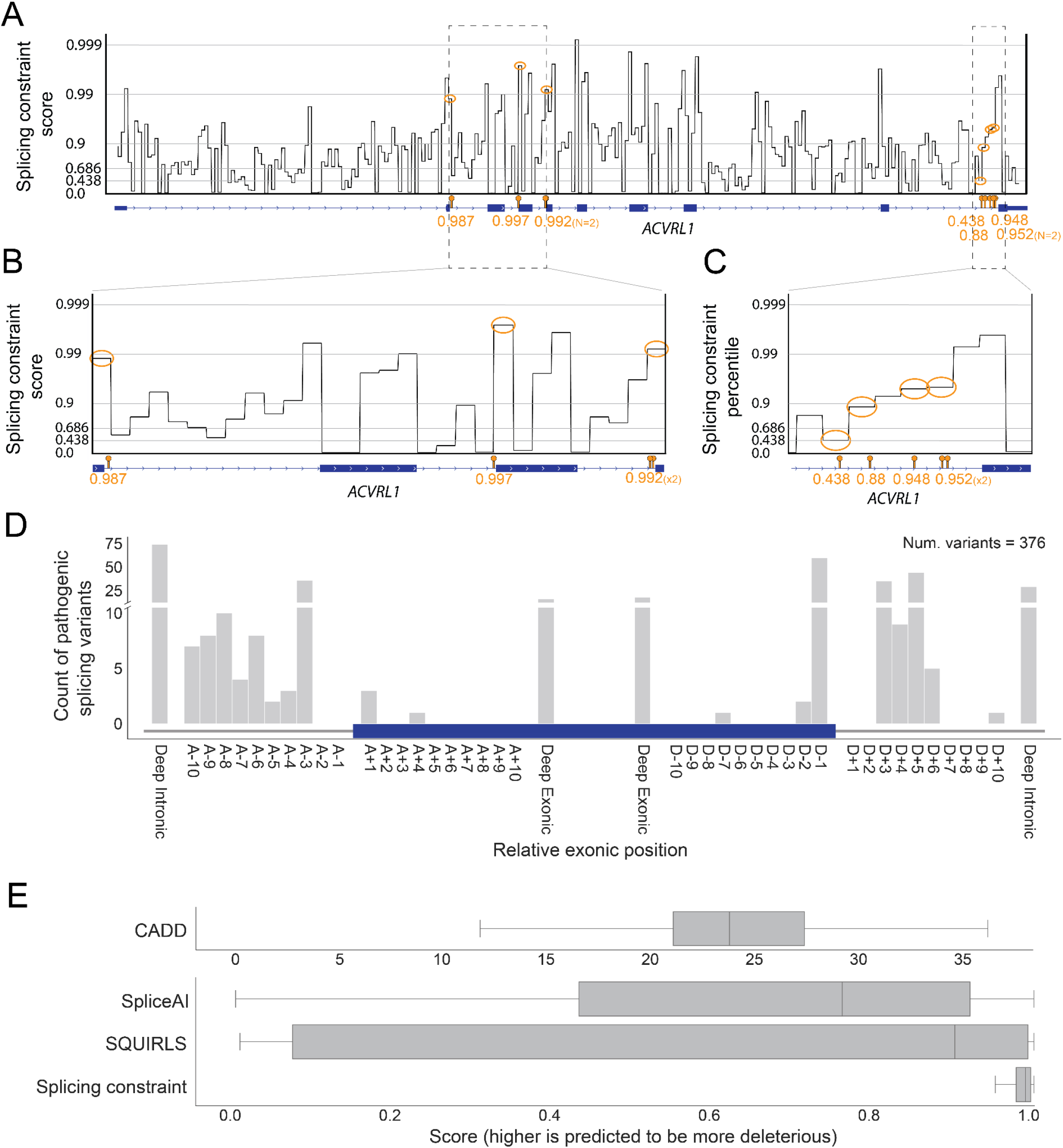
Splicing constraint for pathogenic alternative splicing variants. (**A**) Regional constraint profile for the *ACVRL1* gene with pathogenic splice-altering variants that cause HHT. The pathogenic variants are labeled as orange lollipops on the gene track and the corresponding splicing constraint region is circled in orange on the ConSplice track. The ConSplice score is provided below the variant lollipop. The y-axis depicts the regional constraint score log10 scale. (**B-C**) Zoomed-in view of the genomic regions that harbor the pathogenic variants in ACVRL1 seen in A. (**D**) The distribution of the manually curated pathogenic variants relative to the exonic position. The blue bar represents an exon while the gray bars represent introns around the exon. Oriented 5’ to 3’ from left to right, with the acceptor side on the left and the donor side on the right. Any variant > 10bp away from the exon-intron junction in the intron or exon are labeled “Deep Intronic”, or “Deep Exonic”, respectively. Canonical acceptor and donor sites = A-1, A-2., D+1, D+2. (**E**) The distribution of scores assigned by each method to the pathogenic variants in the set. The IQR of the boxes ranges from the 25th to 75th percentile. The vertical black line in each box represents the median score for that method. The whiskers are 1.5X the IQR. Outliers are not plotted.

We excluded variants at canonical donor and acceptor splice sites from this list of pathogenic splicing variants to evaluate the performance of the splicing constraint model outside of regions that are commonly used when prioritizing variants in rare disease clinical diagnosis (**Figure 4D**). We also included SpliceAI^76^, SQUIRLS^77^, and CADD v1.6^124^ to compare our performance to state-of-the-art splicing prediction and interpretation methods. We excluded other splicing prediction software because they cannot score all of the deep intronic variants in the set. Furthermore, many tools have been previously compared to SpliceAI, SQUIRLS, or CADD. Although SpliceAI, SQUIRLS, and CADD perform well, the regional constraint model significantly scores these variants much higher than each of these tools (**Figure 4E**).

Frésard et al. recently used RNA-seq to identify a pathogenic splicing mutation around the 3’ end of exon 5 in *ASAH1* that leads to exon 6 skipping and causes Spinal Muscular Atrophy with Progressive Myoclonic Epilepsy (SMAPME [MIM: 159950]).^125^ Motivated by this finding, we evaluated the splicing constraint model’s ability to isolate such deleterious splicing events in the absence of RNA sequencing (RNA-seq) data. We find that it accurately identifies the splicing constraint of this region with a score of 0.989 (**Supplemental Figure S13A**). Murdock et al.^126^ used RNA-seq to identify deleterious changes in splicing and expression in a cohort of undiagnosed patients from the Undiagnosed Disease Network (UDN). They identified multiple deleterious splice-altering variants that explained disease pathology. The variants identified included: a deep intronic pathogenic splicing variant in *PQBP1* that leads to an out-of-frame pseudoexon and causes X-linked recessive Renpenning Syndrome (RENS1 [MIM: 309500]); a deep intronic pathogenic splicing variant in *HNRNPK* that resulting in intron retention that caused Au-Kline Syndrome (AUKS [MIM: 616580]); and an intronic pathogenic splicing variant in *RPL13* that lead to intron retention and causes Spondyloepimetaphyseal Dysplasia (SEMD [MIM: 618728]). Similarly, the splicing constraint model scored each region above 0.98 (**Supplemental Figure 13B-D**).

### Variant interpretation with ConSpliceML

While regional constraint identifies genomic regions harboring nucleotides with the potential for deleterious splicing when mutated, not all loci in each region are constrained. Consequently, solely using a region’s constraint to prioritize individual residues will result in false-positive predictions of deleterious splicing. Therefore, we created ConSpliceML to provide a more accurate method for prioritizing individual genetic variants. ConSpliceML uses a Random Forest (RF) classifier to combine regional splicing constraint measures with per-nucleotide alternative splicing predictions from SpliceAI and SQUIRLS (**Methods**). We chose to use both SpliceAI and SQUIRLS since they each perform well at predicting alternative splicing,^77^ yet also capture variants missed by the other.

To train and evaluate ConSpliceML, we used the Human Gene Mutation Database (HGMD),^127–129^ to collect 18,317 disease-causing, alternative splicing variants. In addition, we collected 48,978 benign variants from a collection of de novo mutations (DNMs) identified in whole-genome sequencing of multi-generational^130^ and two generation^131^ families from Utah and Iceland, respectively, as well as validated benign alternative splicing variants from GTEx curated by Jaganathan et al.^76^ (**Supplemental Table S5**). We only included variants in protein-coding genes with SpliceAI, SQUIRLS, CADD, and regional splicing constraint scores. The final set represents 95% of the autosomal pathogenic variants and 36% of autosomal benign variants after filtering. Many of the benign variants are removed because they do not fall within protein-coding genes. We split this variant collection into a training set and test set using 60% of the variants for training and 40% for testing. To further avoid overfitting, we exlcuded variants in the training set if a variant in the test set was found in the same splicing constraint region. Furthermore, we performed training and validation using five-fold cross-validation with the training set to assess training accuracy performance and training consistency across different folds (**Supplemental Figure S14**). To comp are the performance between CADD, SpliceAI, SQUIRLS, and ConSpliceML, we trained the ConSpliceML model using the 60% training set and tested the performance of each method using the 40% test set.

We analyzed each method’s precision and recall for differentiating pathogenic and benign splice-altering variants at various thresholds. With all the variants in the test set, we find that each method performs well, with ConSpliceML performing the best with an average precision of 0.97 and SpliceAI as the second best with an average precision of 0.96 (**Supplemental Figure S15**). However, since variants at canonical donor and acceptor splicing sites are routinely prioritized without the need for annotations during rare disease clinical diagnosis, we evaluated how well each method could isolate deleterious splicing variants outside of the canonical splice sites. We find a moderate decrease in performance for each method, with ConSpliceML continuing to outperform all other methods (**Figure 5A**).

**Figure 5:**
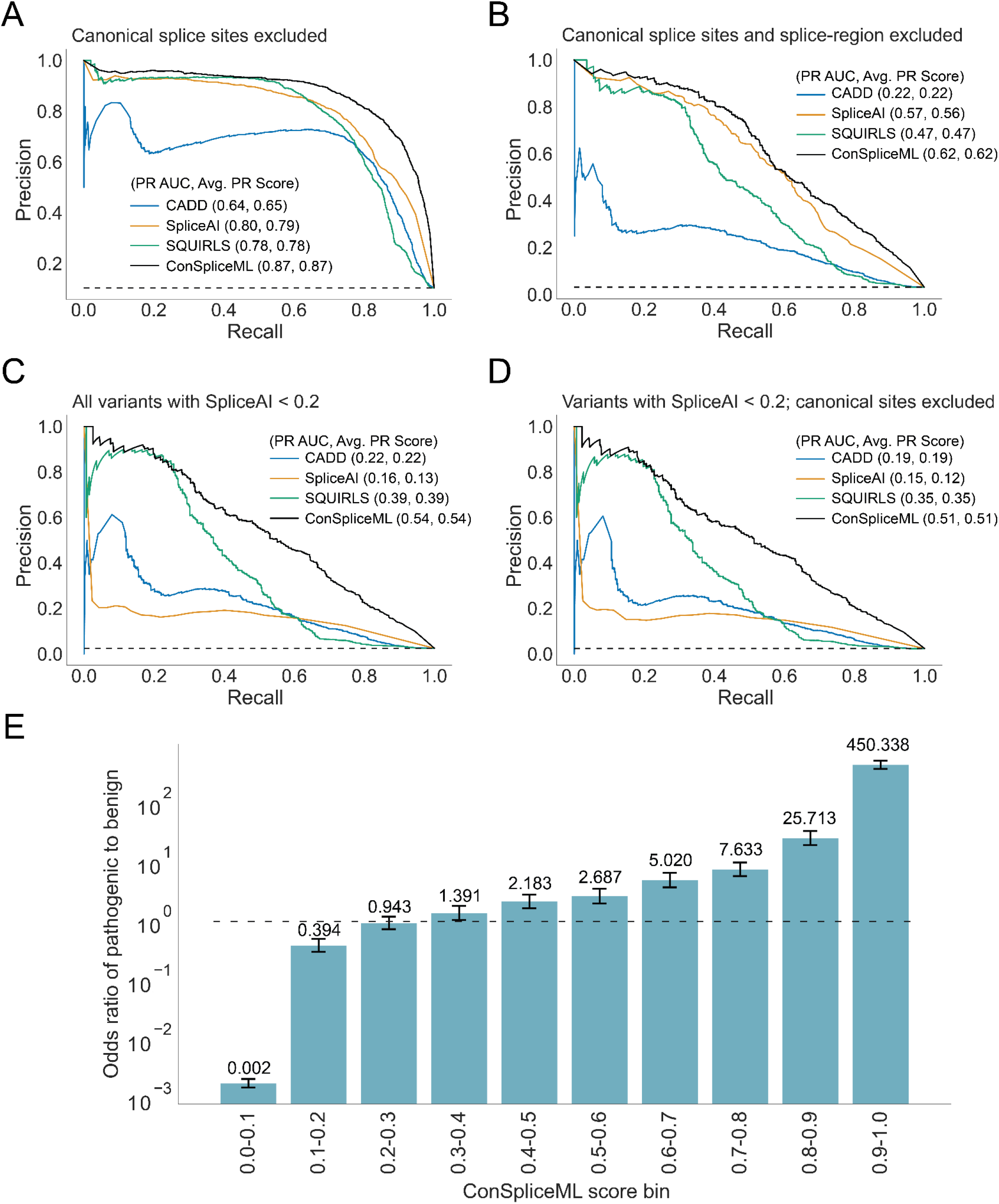
Splicing prediction and interpretation performance. (**A-D**) Precision-Recall (PR) curves depicting the performance of CADD, SpliceAI, SQUIRLS, and ConSpliceML in differentiating pathogenic splice-altering variants from benign variants. The dotted horizontal line represents the baseline value, meaning a random chance prediction. The Precision-Recall Area Under the Curve (PR AUC) for each metric is given in the plot legend, along with a more accurate measurement of performance called the Average Precision Score (Avg. PR Score). (**A**) PR Curve for all pathogenic and benign variants in the test set, excluding the canonical donor and acceptor splice sites. 2,273 pathogenic variants. 19,499 benign variants. (**B**) PR Curve for all pathogenic and benign variants in the test set, excluding variants in the splice region. 627 pathogenic variants. 19,278 benign variants. (**C**) All variants in the filtered test set with a SpliceAI score between 0.0 and 0.2. 477 pathogenic variants. 18,688 benign variants. (**D**) Same as C but excluding variants at the canonical acceptor and donor splice sites. 452 pathogenic variants. 18,686 benign variants. (**E**) The odds ratio enrichment of pathogenic vs. benign variants across the ConSpliceML deciles uses all test set variants. Odds ratio values are highlighted in black above each bar. Error bars represent the 95% CI. The dotted horizontal line represents an odds ratio at 1.

Furthermore, roughly 50% of rare, monogenic disease cases are diagnosed when prioritizing variants in coding sequence, canonical splice sites, or variants within the “splice region” defined by VEP^132^ as eight bases into the intron and three bases into the exon at exon-intron junctions. We, therefore, measured the performance of each method outside the canonical splice sites and “splice regions” commonly used to prioritize variants in rare disease studies. Not surprisingly, we observed a decrease in performance by all methods, with CADD having the greatest drop in performance and ConSpliceML outperforming all other methods (**Figure 5B**).

The authors of SpliceAI recommend prioritizing variants as potentially splice-altering if the SpliceAI score is greater than 0.2.^76,133–135^ However, some pathogenic splicing variants fall below this threshold and, as a result, would be ignored in rare disease analyses. When evaluating performance for variants having a SpliceAI score less than 0.2, we observe that ConSpliceML outperforms all other methods when either canonical splice variants are included or not (**Figure 5C, D**). We excluded the analysis of variants outside the splice region due to an insufficient number of pathogenic variants for evaluation. Although it is not clear why SpliceAI is underperforming for these variants, there is evidence that SpliceAI misses certain alternative splicing variants. For example, Chen et al. found that SpliceAI was unable to fully distinguish between deleterious or neutral splice-altering variants and non-splice-affecting variants at the +2 position of splice donor sites.^136^

Lastly, we measured the enrichment of pathogenic versus benign variants observed across ConSpliceML score ranges to establish a rational scoring threshold for rare disease analyses (**Figure 5E, Supplemental Figure S16**). We observed a significant difference in the enrichment scores between pathogenic and benign variants starting at a ConSpliceML score of 0.5 (**Figure 5E**). Further restricting variants to those outside of canonical sites and the splice region revealed a significant enrichment starting at a ConSpliceML score of 0.3 and 0.2, respectively (**Supplemental Figure S16A, B**). Therefore, we recommend that when using ConSpliceML, a score of 0.5 be used as a preliminary cutoff to distinguish pathogenic and benign splice altering variants. Subsequent iteration could reduce the cutoff to a lower ConSpliceML score to improve the sensitivity of identifying deleterious deep intronic and exonic splicing variants outside the splice region.

## Discussion

Splicing is a highly regulated cellular process that, when disrupted, can lead to disease. Beyond a few nucleotides around exon-intron junctions, variants that could yield aberrant splicing are typically ignored during clinical diagnosis. There is growing evidence that cryptic splicing variants may underlie unsolved cases of rare disease and is a factor in the low diagnostic yield. Although multiple tools have been developed to improve the identification and interpretation of splice-altering variants, they largely underperform when interpreting intronic and exonic splice-altering variants outside the splice region. We developed a model of genetic constraint against aberrant splicing that combines patterns of genetic variation observed in a large cohort of ostensibly healthy individuals with per-nucleotide splicing predictions from SpliceAI. To our knowledge, this model is the first constraint metric built specifically for aberrant splicing that can score the entire gene body rather than solely loci near coding regions.

However, since our model measures splicing constraint in genomic regions, it is insufficient to provide an accurate score at the nucleotide level for variant interpretation. Therefore, we created ConSpliceML, which combines regional constraint with per-nucleotide splicing predictions from SpliceAI and SQUIRLS, providing a per-nucleotide prediction of pathogenic splicing in protein-coding genes. ConSpliceML outperforms state-of-the-art splicing prediction methods at differentiating pathogenic and benign splice altering variants, demonstrating the impact of including constraint on measures of potential deleteriousness.

However, ConSpliceML has notable limitations. First, population datasets, such as gnomAD, are far from reaching saturation for splicing variation, a common problem for pLoF variants.^80^ This limitation will lessen as the number of human genomes increases in population genomic datasets. Another benefit of increasing the number of individuals in the human population datasets is that it provides additional power to increase the resolution of regional constraint models, which will help identify more focalized regions in a gene driving the splicing constraint. Secondly, there is a limited number of available pathogenic splicing variants, especially in deep intronic and exonic regions, in our truth set. This small set of variants limits our ability to measure ConSpliceML’s performance outside of canonical splicing regions. Further evaluation of ConSpliceML and other methods will be necessary as the community identifies additional deep intronic and exonic splicing variants in studies of rare human diseases.

## Conclusion

Genetic constraint facilitates the interpretation of genomic variants in rare disease studies. ConSpliceML outperforms other prediction and interpretation methods for splicing variants at the canonical donor and acceptor sites, in the splice region, and in deep intronic and exonic regions of protein-coding genes. We provide a command-line tool (https://github.com/mikecormier/ConSplice) that generates the splicing constraint model, trains ConSpliceML, and scores variants with both splicing constraint and ConSpliceML. We also offer a VCF^137^ file with precomputed regional constraint score and ConSpliceML scores for all possible SNVs that could occur in protein-coding genes. We suggest an initial pathogenic cutoff of 0.5 when using ConSpliceML and subsequently reducing the cutoff for additional iterations to improve the sensitivity to identify deleterious splicing variants outside of the canonical splice regions. These resources allow for easy distribution of ConSpliceML to the research and clinical communities for improved splicing variant detection and interpretation. We anticipate that ConSpliceML will facilitate the clinical diagnoses for patients with rare diseases, as it identifies missing pathogenic variants driving disease pathology.

## Methods

### Estimating substitution probabilities

Observed variants used to construct the substitution probabilities were identified using the genetic variation found in 76,156 individuals from v3.1.1 of gnomAD.^80^ Variants were used only if they passed the following filtering criteria: the variant was a single nucleotide variant (SNV), had a PASS filter, had an allele count >=1, had a least 10X coverage in 50% of the samples, had a reference allele that matched the reference genome, was in a protein-coding gene determined by GENCODE^138,139^ v34 transcripts, had an associated SpliceAI score prediction, and was not in the pathogenic or benign variant truth set. Additionally, any self-chain or segmental duplication region with a 95% or more identity to another region of the genome was excluded from the splicing constraint model. Because of the difference in constraint between the autosome and the X chromosome, two separate substitution probability matrices and two separate splicing constraint models were generated representing the autosome and the X chromosome. The Y chromosome is excluded in these models.

Relative substitution probabilities were calculated by creating a substitution matrix that modeled each reference allele changing (or not) to any of the other alleles across all protein-coding genes. The reference to nonreference cells in the 4×4 matrix track the number of per-reference to alternative observed variants from gnomAD that pass the filters mentioned above, while the reference to reference cells tracked the number of sites in protein-coding genes that did not have a gnomAD variant. A reference-specific substitution probability was then calculated using the marginal distribution of counts for each reference allele, where the probability represents the normalized counts of reference to non-reference alleles, which is the normalized difference between the total number of sites and the number of sites with no gnomAD variant for that reference allele.

These substitution probabilities were further refined using SpliceAI^76^ predictions, which provide an alternative splicing prediction for each nucleotide considered. SpliceAI allows us to consider the splicing specific substitution rate in coding and non-coding regions of genes where before only the canonical donor and acceptor splice sites could confidently be used. Thus, the substitution probabilities can be transformed into splicing aware expectations, utilizing every site in protein-coding genes. Furthermore, SpliceAI uses the local sequence context around a nucleotide, 5kb upstream and 5kb downstream, to predict the effect of each nucleotide on splicing, thus adding a broader sequence context to the expectation calculation. SpliceAI provides four predictions for each nucleotide, including Acceptor Gain (AG), Acceptor Loss (AL), Donor Gain (DG), and Donor Loss (DL). We chose to use the sum of the four SpliceAI scores, sum(AG, AL, DG, DL), at a single site to account for nucleotides that may contribute to more than one of the four alternative splicing scenarios based on sequence context. For a given reference allele, SpliceAI provides predictions for each possible alternative SNV allele at that site. For sites without a gnomAD variant that passed the filters mentioned above, the sum of the four SpliceAI scores for each reference to alternative allele at that position was calculated and the max sum SpliceAI score was selected. For sites with a variant, the sum SpliceAI score was calculated using the four SpliceAI scores for the variant’s specific reference to the alternative allele.

The substitution matrix was updated by separating the reference to alternative allele counts into eight SpliceAI score range bins (**Supplemental Figure S1A**) for each of the four reference alleles changing (or not) to the four alternative alleles. The final substitution probabilities were calculated by merging all GENCODE protein-coding transcripts, separated by positive and negative strand, and counting the number of gnomAD variants or sites without gnomAD variants across the merged transcripts, separated by SpliceAI score prediction and reference allele to alternative allele combination. The substitution probabilities were calculated as mentioned above, which resulted in 32 substitution probabilities based on a reference allele and SpliceAI score combination (**Figure 1A, Supplemental Figure S1B, Supplemental Table S1,2)**. These substitution probabilities make up the global per-site expectation of seeing an alternative splicing variant based on the reference allele and SpliceAI score prediction for each nucleotide. Canonical GENCODE transcripts, segmental duplication annotations, and self-chain annotations were obtained from the Go Get Data^140^ (GGD) data management system.

### Building the regional model of splicing constraint

To calculate the genetic constraint against aberrant splicing in coding and non-coding regions of the genome, we sought to measure the degree to which the variation observed deviated from what would be expected given the relative substitution probability matrix. Observed counts for a region were measured by adding up the number of gnomAD variants that passed previously stated filters and separated into 32 bins based on each variant’s reference allele and SpliceAI score prediction. Expected counts for the same region were based on the substitution probabilities for each nucleotide in the region, where each nucleotide was assigned an expectation from the substitution probability matrix based on the nucleotides reference allele and SpliceAI prediction, again separated into the 32 bins. Once observed and expected counts were collected for a region, the window was slid down the genome, and the observed and expected counts for the new region were collected. Finally, an Observed / Expected (O/E) ratio was calculated for each region using the region’s observed counts and expected counts. First, an O/E ratio was calculated for each of the 32 bins in a region separately, weighted by the likelihood of that bin contributing to splicing. We include a likelihood weight here to boost the signal for constraint against splicing in a given region and reduce the noise from irrelevant alleles in that same region. Each bin’s likelihood weight was based on the proportion of sites in the genome with that reference allele and SpliceAI score range combination, where the weight was equal to one over the genome-wide proportion for that bin. Therefore, irrelevant alleles that do not contribute to splicing will be down-weighted, reducing noise, while relevant alleles that do contribute to splicing will be up-weighted, boosting the signal. Second, weighted O/E ratios for all 32 bins for a single region were summed together to get the final O/E ratio for that region.

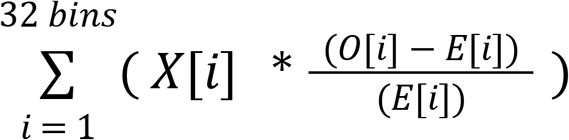

- <u>32 bins</u> = The 32 bins based on the combination of reference alleles and SpliceAI score ranges.
- X = A weight based on the likelihood that bin *i* contributes to splicing (See below).
- <u>O</u> = The observed counts at a specific reference allele + SpliceAI score range bin *i.*
- <u>E</u> = The expected counts at a specific reference allele + SpliceAI score range bin *i.*

To normalize the observed and expected counts across each of the 32 bins per region, we set the observed and expected counts to a default value of 1.0. Using the equation above, these default values lead to a regional O/E distribution being centered at 0 rather than at 1. Additionally, these default values with the O/E equation above remove the lower limit of 0 seen in traditional O/E and chi-squared statistics to a continuous negative value lower limit, allowing for higher constraint to be represented by a larger negative value with a null O/E value expectation of 0.0, and higher O/E values representing an increase in constraint. The mean O/E value for single exon genes was arbitrarily increased in the positive direction to account for the absence of splicing in single-exon genes.

Once an O/E score was calculated for every region, the O/E scores were sorted from largest to smallest and assigned a normalized score between 0.0 and 1.0. A constraint score of 1.0 represents the most constrained region to splicing, while a constraint score of 0.0 represents the most tolerant region to splicing.

Due to the difference between the autosome and the X chromosome, substitution probabilities and splicing constraint models were generated separately for both the autosome and the X chromosome. The pseudoautosomal regions (PAR) on the X chromosome were excluded from the X chromosome model. (GRCh38) PAR 1 = chrX:10000-2781479, PAR 2 = chrX:155701382-156030895.

### Evaluation of likelihood weights and optimal region sizes

To determine the best weighting approach among the six different likelihood weights used to develop the constraint model, performance was compared across each of the different weights using the pathogenic variants from HGMD and the validated benign alternative splicing variants in the benign set from GTEx (see “Pathogenic and benign variant truth set” methods section below). We found that the regional constraint model performed best when using the 1 / Proportion weight (**Supplemental Figure S17-19**). **Supplemental Figure S17-19** also shows that the linear weighted model performs better for larger regions.

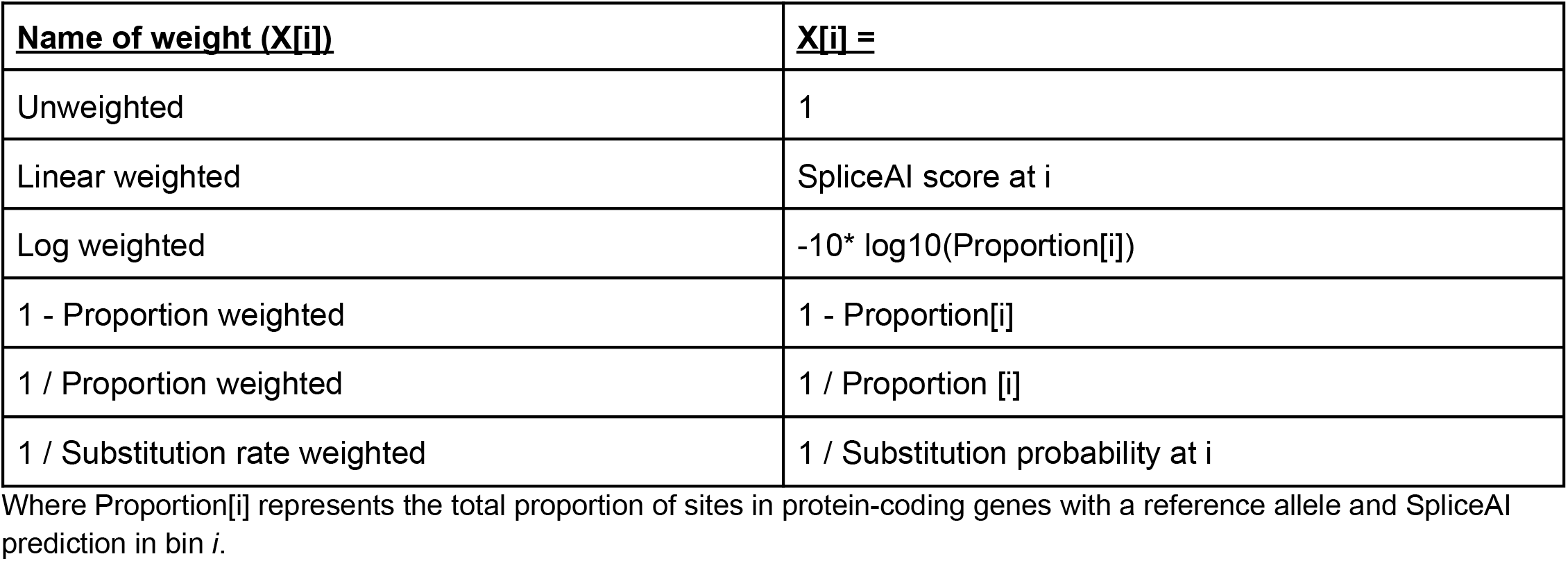

| Name of weight (X[i]) | X[i] = |
| --- | --- |
| Unweighted | 1 |
| Linear weighted | SpliceAI score at i |
| Log weighted | $-10 * \log_{10}(\text{Proportion}[i])$ |
| 1 - Proportion weighted | 1 - Proportion[i] |
| 1 / Proportion weighted | 1 / Proportion [i] |
| 1 / Substitution rate weighted | 1 / Substitution probability at i |

Where Proportion[i] represents the total proportion of sites in protein-coding genes with a reference allele and SpliceAI prediction in bin *i*.

Both the genic and regional constraint models were developed using a window-based approach. The gene size defined the window sizes for the gene model. Intragenic regional models were defined by a window either equal to or smaller than the size of a gene. Specific regions were excluded from the model if there was less than 80% of the nucleotides in that region with a SpliceAI score. To maintain a sufficiently large window to calculate constraint, we only included windows that were above the significant cutoff for informative sites based on the critical value determined using the F distribution. The number of informative sites by window size was modeled with a Poisson distribution where lambda was set to the genome-wide proportion of sites in protein-coding genes that were predicted to affect splicing by SpliceAI with a SpliceAI score > 0.0 (**Supplemental Figure S1A, S7**). The significance cutoff was based on the F distribution at an alpha value of 0.025 using 7 degrees of freedom representing the 8 proportion bins used to calculate the proportion of informative sites. The critical value from the F distribution table is 6.5415, meaning a window needs at least 6.5415 informative variants to be significant. **Supplemental Figure S7** shows the number of informative sites per window size. Based on the significance cutoff, a window size of 50bp or greater was used to generate the constraint model. **Supplemental Figure S10** shows the performance of various windows sizes using the 1 / Proportion weight constraint model, where performance is assessed using the HGMD pathogenic variants and validated splice-altering benign variants from GTEx in the truth set. **Figure 3A** and **Supplemental Figure S8** provide additional testing to evaluate the ability of each window size to capture focal constraint around 5’ and 3’ splice regions.

### Regional splicing constraint score bins

For multiple analyses in this manuscript, the regional constraint scores are separated into decile bins. For each bin, any gene or variant with a regional constraint score within the bin’s range was included in that bin. For all but the last bin, genes and variants were included if their regional constraint score was equal to or greater than the lower value of the bin range and less than the upper value of the range. For the last bin, any gene or variant with a regional constraint score greater than or equal to the lower value of the bin range and less than or equal to the upper value of the range were included in that bin.

### Gene sets used to assess and compare splicing constraint

We used multiple gene sets to evaluate the genic splicing constraint scores based on previously established constraint expectations for those gene sets.

Gene Sets:

- <u>Haploinsufficient genes</u>: We used a set of 294 genes determined to be dosage-sensitive in the ClinGen data set^141^ and hosted on the Macarthur lab gene list GitHub page at https://github.com/macarthur-lab/gene_lists/blob/master/lists/clingen_level3_genes_2018_09_13.tsv.
- <u>Autosomal Dominant genes</u>: We used a combined set of 709 genes that were shown to follow an autosomal dominant inheritance pattern determined by Blekhman et al.^142^ and Berg et al.^143^ hosted on the Macarthur lab GitHub page at: https://github.com/macarthur-lab/gene_lists/blob/master/lists/berg_ad.tsv.
- <u>Autosomal Recessive genes</u>: We used a combined set of 1183 genes that were shown to follow an autosomal recessive inheritance pattern by Blekhman et al.^142^ and Berg et al.^143^ hosted on the Macarthur lab GitHub page at: https://github.com/macarthur-lab/gene_lists/blob/master/lists/all_ar.tsv.
- <u>Olfactory Receptor genes</u>: We used a set of 371 Olfactory Receptor genes from the Mainland et al.^144^ paper hosted on the Macarthur lab GitHub page at: https://github.com/macarthur-lab/gene_lists/blob/master/lists/olfactory_receptors.tsv.
- <u>CRISPR Essential and Nonessential genes</u>: We used a set of 683 essential genes and 913 nonessential genes determined to be essential or nonessential, respectively, by CRISPR screens from Hart et al.^84^ hosted on the Macarthur lab GitHub page at: essential genes: https://github.com/macarthur-lab/gene_lists/blob/master/lists/CEGv2_subset_universe.tsv, nonessential genes: https://github.com/macarthur-lab/gene_lists/blob/master/lists/NEGv1_subset_universe.tsv.
- <u>Homozygous LoF tolerant genes</u>: We used a set of 1815 genes tolerant to high confident homozygous loss of function mutations in at least one individual in gnomAD found in Supplemental Table S7 of the gnomAD paper^80^: https://static-content.springer.com/esm/art%3A10.1038%2Fs41586-020-2308-7/MediaObjects/41586_2020_2308_MOESM4_ESM.zip
- <u>Developmental Delay and Intellectual Disability (DD/ID) genes</u>: We obtained a set of 2072 genes collected across multiple Deciphering Developmental Delay (DDD) studies hosted on the Gene 2 Phenotype^90^ website: https://www.ebi.ac.uk/gene2phenotype/, csv file: https://www.ebi.ac.uk/gene2phenotype/downloads/DDG2P.csv.gz.

Genes in the Haploinsufficient, Autosomal Dominant, Autosomal Recessive, Olfactory Receptor, DD/ID, CRISPR Essential, and CRISPR Nonessential gene sets were each normalized to a value of one across the genic splicing constraint score spectrum to compare the proportion/fraction of genes in each constraint decile bin. Genes in the Homozygous LoF tolerant gene set were separated into constraint decile bins. For each genic splicing constraint decile bin, the number of Homozygous LoF tolerant genes in a bin was divided by the total number of genes in that same bin to get the percent of Homozygous LoF tolerant genes by decile.

### pLI and LOEUF constraint scores

The genic splicing constraint metric was compared to the probability of loss-of-function intolerance (pLI)^81^ and the loss-of-function observed/expected upper bound fraction (LOEUF)^80^ metrics to establish if the splicing constraint metric captured known patterns of constraint. pLI and LOEUF were downloaded from the gnomAD browser using the constraint download link https://gnomad-public-us-east-1.s3.amazonaws.com/legacy/exac_browser/forweb_cleaned_exac_r03_march16_z_data_pLI_CNV-final.txt.gz for pLI and the constraint download link https://gnomad-public-us-east-1.s3.amazonaws.com/release/2.1.1/constraint/gnomad.v2.1.1.lof_metrics.by_gene.txt.bgz for LOEUF. The “pLI” column and the “oe_lof_upper_bin” column were used for the pLI score and the LOEUF score in their respective files.

Splicing constraint and pLI scores are ranked from least to most constrained from 0.0 to 1.0. LOEUF scores are ranked in the opposite direction with the most constrained gene assigned a LEOUF decile of 0 and the least constrained gene assigned a LOEUF decile of 9. To compared LOEUF to pLI and splicing constraint, we reversed the LOEUF deciles so that the ranking was oriented in the same direction as pLI and splicing constraint, and relabeled the deciles to match the pLI and splicing constraint decile bins. Therefore, LOEUF deciles in this mansucript are as follows:

| **<u>Original LOEUF decile</u>** | **<u>Reversed and relabeled LOEUF decile</u>** |
| --- | --- |
| 9 | 0.0-0.1 |
| 8 | 0.1-0.2 |
| 7 | 0.2-0.3 |
| 6 | 0.3-0.4 |
| 5 | 0.4-0.5 |
| 4 | 0.5-0.6 |
| 3 | 0.6-0.7 |
| 2 | 0.7-0.8 |
| 1 | 0.8-0.9 |
| 0 | 0.9-1.0 |

### Comparing genic splicing constraint to gene dosage sensitivity using V^G^ estimates from GTEx

To assess the correlation between splicing constraint and gene dosage, we used the average genetic variation in gene expression (V^G^) values created from GTEx data by Mohammadi et al.^92^ located in their Supplemental Table S1. The average V^G^ values are a per gene weighted harmonic mean across tissues in natural log units. We removed any outlier V^G^ value that was > 10 standard deviations from the mean. We assigned each gene its cross-tissue average V^G^ value and separated them into ConSplice score decile bins. We plotted the distribution of average V^G^ values per constraint decile bin as box plots. The interquartile range (IQR) of the boxes ranges from the 25th to 75th percentile. The horizontal line in each box represents the median. The whiskers are 1.5X the IQR. Outlier values are not included in the plot.

To test for a significant negative correlation between the splicing constraint of a gene and its estimated genic expression, we ran an ordinary least squares (OLS) linear regression using the splicing constraint scores and V^G^ values. We also controlled for the number of exons per gene in the regression to rule out any covariant significance driven by the number of splicing events in a gene. The mean fold difference of variance in gene expression between the lowest and highest constraint decile bins was calculated by taking the mean V^G^ value in the 0.0-0.1 decile and dividing by the mean V^G^ values in the 0.9-1.0 decile. We further tested for a significant difference in V^G^ distributions between different constraint bins. We compared each bin to every other bin in a pair-wise manner using a t-test of independence. The p-values were adjusted using the Benjamini Hochberg approach with an alpha value of 0.01. FDR adjusted p-values can be found in **Supplemental Figure S4**.

### Comparing genic splicing constraint to GTEx splicing outliers

Splicing Outliers from Ferraro et al.^95^ were obtained from the GTEx portal within the version 8 data release file named “gtexV8.sOutlier.stats.globalOutliers.removed.txt.gz” under the “Outlier Calls” heading: https://storage.googleapis.com/gtex_analysis_v8/outlier_calls/GTEx_v8_outlier_calls.zip. The data within the sOutiler file represent the per gene cross-tissue p-values for splicing outliers found for each GTEx sample, where the p-values were determined using the SPOT^95^ metric. For a sOutlier to be included in this dataset for a sample, it needed to have been present in multiple tissues based on the requirements from Ferraro et al. These events represent a highly confident set of sOutliers.

Ferraro et al. determined that a p-value cutoff of 0.0027 was adequate to determine whether or not a cross-tissue splicing outlier was significant or not. We used the same p-value cutoff and designated significant sOutliers as those with a p-value < 0.0027. For each gene, we counted the number of individuals in GTEx that had a significant cross-tissue sOutlier p-value.

Using the gene designation within the sOutlier file, we added the associated constraint scores. We counted the number of genes with 0, 1, 2, 3, 4, 5, 6, 7, 8, 9, 10, and > 10 sOutliers per each constraint decile bin. That is, how many genes within a constraint decile bin have zero significant sOutilers, one significant sOutlier, and so on. We then normalized the total number of sOutlier genes (i.e., genes from the sOutlier file) for each constraint decile bin. The total normalized number of sOultier genes in a constraint decile bin will sum to 1. **Figure 3C** shows the normalized values for genes with zero sOutliers per constraint bin. **Supplemental Figure S5B** shows the normalized values for the genes with at least one sOutlier. Significance was determined using OLS linear regression.

### Assessing regional constraint in constitutive vs cassette Exons

We used constitutive and cassette exons from the HEXEvent database^98^ to evaluate the regional splicing constraint metric around exons. Exon definitions were downloaded from the HEXEvent website http://hexevent.mmg.uci.edu/cgi-bin/HEXEvent/HEXEventWEB.cgi for the GRCh38 genome build. For each exon, HEXEvent provides a measure of exon inclusion called the constitutive level (constLevel). The inclusion level for each exon provided by HEXEvent was used to determine “Full Inclusion” constitutive exons, “High Inclusion” cassette exons, and “Moderate Inclusion” cassette exons. High inclusion cassette exons were defined as exons with an inclusion >= 90% and < 100% based on previous work defining a 90% cutoff.^98,99^ Moderate inclusion cassette exons included all exons with an inclusion rate < 90%.

For each exon, a regional constraint score was identified at 25bp intervals from the 5’ and 3’ ends of the exon, moving upstream and downstream of the exon into the intron, respectively. These intervals continued until the upstream and downstream exon had been reached. The list of regional scores for exons on the negative strand was re-oriented to match the positive strand in a 5’ to 3’ manner. Solid lines represent the median ConSplice score at each 25bp interval. The opaque shading around the solid lines represents the 95% confidence interval calculated using bootstrap with 1000 iterations.

### Splicing unaware model of constraint

To determine if the regional model of splicing constraint was identifying the constraint of coding regions compared to non-coding regions alone or identifying the signal of splicing beyond coding regions, we created multiple regional models of constraint unaware of splicing to compare to the splicing aware models of constraint. The same approach already described for the splicing constraint model was used to create the splicing unaware model of constraint. The difference between the two models lies in the inclusion or exclusion of SpliceAI predictions to inform the model, where the splicing aware model of constraint uses the SpliceAI prediction and the splicing unaware model of constraint excludes the SpliceAI predictions. Additionally, since SpliceAI predictions are excluded from the splicing unaware model of constraint, the splicing likelihood weights used to calculate constraint were also excluded, representing an unweighted model.

### Assessing regional constraint in poison exons

Genomic coordinates for the three poison exons in *SCN1A* seen in **Figure 3B** and **Supplemental Figure S12** that cause nonsense-mediated decay (NMD) leading to Dravet Syndrome (DRVT [MIM: 607208]) were collected from Steward et al.,^101^ (**Supplemental Table S3**). IGV^145^ was used to visualize the constraint score profiles around each exon feature. Constraint scores reflect the 50bp regional constraint model. **Supplemental Table S3** also contains the coordinates of the three randomly selected exons in *SCN1A* seen in **Figure 3C** and **Supplemental Figure S12**. To identify the ranking for each poison exon in *SCN1A,* all constraint scores in *SCN1A* were sorted and assigned a score between 0.0 and 1.0. The max constraint score that overlapped each poison exon was at or above the 92nd percentile for *SCN1A* constraint scores.

Coding sequencing (CDS) annotations were used to distinguish coding and noncoding sequences in *SCN1A.* Canonical splice sites were determined based on the −1, −2, +1, and +2 positions around each CDS exon, excluding the −1 and −2 positions of the first exon and the +1 and +2 positions of the last exon which do not participate in splicing. The distribution of regional constraint scores was plotted based on if the region overlapped a CDS region or not, or a canonical splice site (**Supplemental Figure S11**). IQR of the boxes ranges from the 25th to 75th percentile. The horizontal line in each box represents the median. The whiskers are 1.5X the IQR. The max regional constraint score for each poison exon is plotted as points along the distribution. The max regional constraint score for each poison exon was in the top 2% of scores for noncoding regions in *SCN1A,* and is similar to the constraint in both CDS regions and canonical splice site regions.

### Manually curated pathogenic splice altering variants

We identified 376 autosomal noncanonical pathogenic splice altering variants by reviewing scientific medical literature. Each variant in this list has functional support for its effect on splicing and an association with a Mendelian disorder. Variants at the canonical donor and acceptor splice sites were explicitly excluded from this set. For each variant, we include PubMed ids for the papers the describe the variant, dbSNP^146^ RS ids, Clinvar^147^ ids along with review status, gnomAD v2 and v3 allele frequencies, online mendelian inheritance of man (OMIM)^148^ ids, the relative exonic position, and the number of bases from the nearest exon-intron junction. Each variant also has a SpliceAI score, a SQUIRLS score, a CADD v1.6 score, and a 50bp regional constraint score. (**Supplemental Table S4**)

IGV was used to visualize the regional constraint profiles in *ACVRL1* found in **Figure 4A-C**. The variant position was plotted on the gene track as a red lollipop. The 50bp constraint region containing the variant is circled in red. The regional constraint score for each variant is listed below the variant lollipop in red. The right y-axis shows the regional constraint score. The left y-axis shows the log scaled constraint scores using the following equation: −10 * log10(1 - constraint score). The constraint profile is plotted using the log scaled scores. Variants in **Supplemental Figure S13** were identified from Frésard et al.^125^ and Murdock et al.^126^ and plotted using the same approach for the *ACVRL1* gene described above.

The count of variants from the set at each relative exonic position was plotted in the 5’ to 3’ direction from left to right. Variants greater than 10bp away from the exon-intron junction were labeled as deep intronic and deep exonic variants. The distribution of scores for each variant in the set from CADD-Splice (the splicing version of CADD v1.6), SpliceAI, SQURILS, and regional constraint are plotted as boxplots with the IQR ranging from the 25th to the 75th percentile, median scores represented by the black vertical line in each box, and the whiskers 1.5X the IQR. Outliers beyond the whiskers are not plotted.

The 50bp regional constraint model was used for all analyses in this section.

### Pathogenic and benign variant truth set

The 18,317 filtered variants in the pathogenic truth set were identified from the Human Gene Mutation Database^127–129^ (HGMD). Each pathogenic variant leads to deleterious splicing and was labeled as a disease-causing mutation (DM) by HGMD. Variants were downloaded from Q1 of 2021. Variants not on the autosome and that did not have a CADD score, SpliceAI score, SQUIRLS score, and regional constraint score were excluded from the analysis. The relative exonic position was identified using the combination of the “type” and “location” columns in the dataset. Due to licensing agreements with HGMD, we are not allowed to share these variants publicly. For those who do have an HGMD commercial license, we provide the MySQL command to download the HGMD variants in the “Availability of Data and Materials”, section(WGS below.

The 48,978 filtered variants in the benign truth set come from a combination of de novo mutations (DNM) from whole-genome sequencing of multi-generational^130^ and two generation^131^ families, and a set of validated benign splice altering variants in GTEx.^76^ Second and third-generation DNMs from the multi-generational family study were downloaded from https://github.com/quinlan-lab/ceph-dnm-manuscript. These DNMs were lifted over from GRCh37 to GRCh38. DNMs from 1,548 two-generation trios were downloaded from Supplemental Table 4 of the Jónsson et al. paper.^131^ These DNMs were already in GRCh38. Validated benign splice altering variants seen in 1-4 individuals in the GTEx RNA-seq data were downloaded from Jaganathan et al.^76^ Variants were lifted over from GRCH37 to GRCh38.

Variants in the benign truth set outside of protein-coding genes were filtered out of the set. All remaining variants were then scored with CADD, SpliceAI, SQUIRLS, and regional constraint. Any variant that did not have a score from CADD, SpliceAI, SQUIRLS, and regional constraint was removed from the set. The relative exonic position was added to each variant using the canonical transcripts in GENCODE v34. This benign set is available in **Supplemental Table S5**.

### The ConSpliceML model

The ConSpliceML model is deployed using a Random Forest (RF) ensemble machine learning approach, a supervised classification machine learning algorithm based on a collection of decision trees, implemented using the scikit-learn^149^ python API. The RF uses the regional constraint, SpliceAI, and SQUIRLS scores as a feature vector with a pathogenic or benign label for each variant to create the decision trees that make up the model. ConSpliceML is made up of 1000 decision trees. The ConSpliceML model provides a prediction at a per-base resolution on a range from 0.0 to 1.0, describing the probability that a variant is a pathogenic splice-altering variant. The model used in this manuscript can be found in the manuscript GitHub repository at https://github.com/mikecormier/ConSplice-manuscript and is implemented as a python module found at https://github.com/mikecormier/ConSplice. The precomputed ConSpliceML scores in the vcf file we provide were generated using the ConSpliceML python module at https://github.com/mikecormier/ConSplice, where the model was trained using the full collection of pathogenic and benign variants in the truth set.

### Scoring Variants

We used CADD v1.6, also known as the splicing version of CADD called CADD-Splice, for the CADD scores used in our dataset. GRCh38 CADD v1.6 scores for all possible SNVs were downloaded from the CADD website at https://cadd.gs.washington.edu/download. Variants were assigned a CADD score based on the genomic position, reference allele, and alternative allele.

GRCh38 SpliceAI v1.3.1 pre-computed scores for SNVs in protein-coding genes were downloaded following the instructions found on SpliceAI’s GitHub page at https://github.com/Illumina/SpliceAI. Variants were assigned a SpliceAI score using the genomic position, reference allele, alternative allele, and gene name.

SQUIRLS version 1.0.0 was downloaded following the instructions outlined in the SQUILRS documentation at https://squirls.readthedocs.io/en/latest/. Variants were scored using the GRCh38 SQUIRLS CLI.

The regional splicing constraint score was assigned to each variant using the ConSplice CLI: https://github.com/mikecormier/ConSplice.

ConSpliceML scores were generated using the trained RF with the ConSplice, Max SpliceAI, and Max SQUIRLS scores from each test variant as the input feature vector. ConSpliceML then provides a pathogenic probability score that ranges from 0.0 to 1.0.

### Training and Testing ConSpliceML

We used 60% of the truth set for training the ConSpliceML model and 40% for testing. The test set was excluded from all training of the ConSpliceML model and represents an independent set of variants for testing. We eliminated possible overfitting based on variants in the training and test set not falling in the same splicing constraint region. The variants within a region that containes multiple variants in the truth set were randomly assigned to either the training set or tests set using the 60/40% split. This approach eliminates any variants in the training set falling in the same splicing constraint region as those in the test set. With the training set, we used a stratified five-fold cross-validation approach to train and validate the training accuracy of the ConSpliceML model. Each of the five folds had a distinct 20% of pathogenic and benign variants not shared by any of the other folds used for validation. The variants used to train each fold used the 80% remaining variants in the training set not found in the validation set for that fold. The number of pathogenic and benign variants in each fold was consistent across all folds. Pathogenic and benign variants were labeled as such. Due to a large difference in the number of pathogenic to benign variants, Precision-Recall (PR) was assessed, as it is more suitable for imbalanced datasets. PR curves were generated and compared across each of the five folds to evaluate the training accuracy, stability, and consistency of the ConSpliceML model (**Supplemental Figure S14)**. The combined average performance across all five folds was also assessed in the same plot.

Finally, the ConSpliceML model was trained using the training set, and the performance of CADD, SpliceAI, SQUIRLS, and ConSpliceML was assessed using the hold-out test set. The area under the precision-recall curve (PR AUR) and the average precision score (Ave. PR Score) are provided for each PR curve. The Ave. PR Score is a weighted mean of precision occurring at each threshold weighted by the difference in recall from the previous threshold and is a more accurate measure of performance.

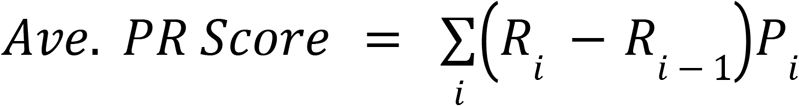

- P_i_ = The precision at the ith threshold.
- R_i_ = The recall at the ith threshold.

The dotted horizontal baseline in each PR plot represents the random chance prediction and is calculated as follows:

- P = The number of pathogenic variants
- B = the number of benign variants

### Odds Ratio Enrichment

To measure the enrichment of pathogenic to benign variants along the ConSpliceML score range, we measure the ratio of pathogenic variants to benign variants in a ConSpliceML decile divided by the ratio of pathogenic to benign variants not in the decile. That is, we calculate the odds ratio (OR) of pathogenic to benign variants at each ConSpliceML decile.

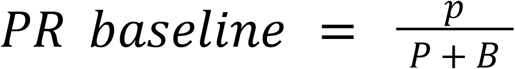

- OR_i_ = The odds ratio of pathogenic to benign variants for the ConSpliceML decile i.
- A_i_ = The number of pathogenic variants in the ConSpliceML decile i.
- B_i_ = The number of benign variants in the ConSpliceML decile i.
- C_not i_ = The number of pathogenic variants not in the ConSpliceML decile i.
- D_not i_ = The number of benign variants not in the ConSpliceML decile i.

We calculate the 95% confidence interval (CI) for each OR as follows:

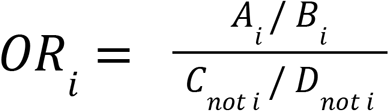

OR by decile was plotted as bar plots with whiskers as the 95% CI. The OR values for each decile are indicated above each bar. The dotted line at an OR of 1 indicates no difference in the ratio of pathogenic to benign variants.

## Abbreviations

ACMG: American College of Medical Genetics
AD genes: Autosomal Dominant genes
AG: Acceptor Gain
AL: Acceptor Loss
AR genes: Autosomal Recessive genes
AUKS: Au-Kline Syndrome
Avg. PR Score: Average Precision Score
CDS: Coding sequencing
Cl: Confidence lnterval
CLl: Command Line Interface
ConSplice: Constrained Splicing
constLevel: Constitutive Level
DD/ID: Developmental Delay and Intellectual Disability
DG: Donor Gain
DL: Donor Loss
DM: Disease-causing Mutations
DNA-seq: DNA sequencing
DNMs: de novo mutations
DRVT: Dravet Syndrome
FDR: False Discovery Rate
GGD: Go Get Data
gnomAD: Genome Aggregate Database
GTEx: Genotype-Tissue Expression
HI genes: Haploinsufficient genes
HGMD: Human Gene Mutation Database
HTT2: Hereditary Hemorrhagic Telangiectasia
IQR: Interquartile range
LOEUF: loss-of-function observed/expected upper-bound fraction
MFS: Marfan Syndrome
NF1: Neurofibromatosis type 1
NMD: Nonsense-Mediated Decay
O/E: Observed over Expected
OLS: Ordinary Least Squares
OPA1: Optic Atrophy 1
OMIM or MIM: Online Mendelian Inheritance of Man
OR: Odds Ratio
OR genes: Olfactory Receptor genes
PAR: Pseudoautosomal Region
pLI: probability of loss-of-function intolerance
pLoF: putative loss-of-function
PR: Precision-Recall
PR AUC: Precision-Recall Area Under the Curve
RB1: Retinoblastoma
RENS1: Renpenning Syndrome
RF: Random Forest
RNA-seq: RNA sequencing
SEMD: Spondyloepimetaphyseal Dysplasia
SMAPME: Spinal Muscular Atrophy with Progressive Myoclonic Epilepsy
SNV: Single Nucleotide Variant
sOutliers: Splicing Outlier
STL1: Stickler Syndrome
UDN: Undiagnosed Disease Network
V^G^: expected genetic variation in gene expression

## Ethics Declaration

### Ethics approval and consent to participate

Not applicable

### Consent for publication

Not applicable

## Availability of Data and Materials

The ConSplice CLI, including a module for splicing constraint and a module for ConSpliceML, is an open-source and publicly available project under an MIT license.

The ConSplice CLI can be found at https://github.com/mikecormier/ConSplice and contains the code necessary to recreate the regional constraint model. Additionally, it contains information on the data required to create the ConSplice model along with how to download and use the data.

The GRCh38 gnomAD80 v3.1.1 vcf file for each chromosome and the per-base sequencing coverage file were downloaded from the gnomAD browser website at https://gnomad.broadinstitute.org/downloads. The separate chromosome vcf files were merged together into one file, decomposed, and normalized. Any gnomAD variant in the truth set was removed from the final gnomAD vcf file to ensure that the splicing constraint model was not trained and tested on the same variants.

The precomputed GRCh38 SpliceAI^76^ v1.3.1 raw SNV vcf files were downloaded from the illumina basespace portal at https://basespace.illumina.com/s/otSPW8hnhaZR following the information available on the SpliceAI GitHub repo at https://github.com/Illumina/SpliceAI.

The GRCh38 gene features from GENCODE^138,139^ v34 used to create the splicing constraint model were downloaded using the GGD140 CLI under the GGD data package name “grch38-canonical-transcript-features-gencode-v1”. (https://gogetdata.github.io/recipes/genomics/Homo_sapiens/GRCh38/grch38-canonical-transcript-features-gencode-v1/README.html).

The segmental duplications and self-chain genomic repeat bed files used to create the splicing constraint model were downloaded using the GGD cli under the GGD data package names “grch38-segmental-dups-ucsc-v1” and “grch38-self-chain-ucsc-v1”, respectively. (https://gogetdata.github.io/recipes/genomics/Homo_sapiens/GRCh38/grch38-segmental-dups-ucsc-v1/README.html, https://gogetdata.github.io/recipes/genomics/Homo_sapiens/GRCh38/grch38-self-chain-ucsc-v1/README.html).

The GGD140 documentation can be found at https://gogetdata.github.io/.

Gene sets used to test the genic splicing constraint model were obtained from the following resources:

- Gene 2 Phenotype^90^: https://github.com/macarthur-lab/gene_lists/blob/master/lists/homozygous_lof_tolerant_twohit.tsv
- Haploinsufficient Genes^141^: https://github.com/macarthur-lab/gene_lists/blob/master/lists/clingen_level3_genes_2018_09_13.tsv.
- Autosomal Dominant Genes^142,143^: https://github.com/macarthur-lab/gene_lists/blob/master/lists/berg_ad.tsv
- Autosomal Recessive Genes^142,143^: https://github.com/macarthur-lab/gene_lists/blob/master/lists/all_ar.tsv
- Olfactory Receptor Genes^144^: https://github.com/macarthur-lab/gene_lists/blob/master/lists/olfactory_receptors.tsv
- CRISPR Essential Genes^84^: https://github.com/macarthur-lab/gene_lists/blob/master/lists/CEGv2_subset_universe.tsv
- CRISPR Non-essential Genes^84^: https://github.com/macarthur-lab/gene_lists/blob/master/lists/NEGv1_subset_universe.tsv
- Homozygous LoF Genes from supplemental Table S7 of the gnomAD paper^80^: https://static-content.springer.com/esm/art%3A10.1038%2Fs41586-020-2308-7/MediaObjects/41586_2020_2308_MOESM4_ESM.zip

V^G^ values created from GTEx data were downloaded from Supplemental Table S1 in the Mohammadi et al.^92^ paper.

Splicing Outliers from Ferraro et al.^95^ were obtained from the GTEx portal https://www.gtexportal.org/home/ within the version 8 data release file named “gtexV8.sOutlier.stats.globalOutliers.removed.txt.gz” under the “Outlier Calls” heading: https://storage.googleapis.com/gtex_analysis_v8/outlier_calls/GTEx_v8_outlier_calls.zip.

The GRCh38 constitutive and cassette exons were downloaded from the HEXEvent website http://hexevent.mmg.uci.edu/cgi-bin/HEXEvent/HEXEventWEB.cgi.

The manually curated noncanonical pathogenic splicing variants are available in **Supplemental Table S4** of this manuscript.

SQUIRLS v1.0.0 was downloaded following the instructions outlined in the SQUIRLS documentation: https://squirls.readthedocs.io/en/latest/.

The disease-causing, splice-altering variants from HGMD were downloaded from the HGMD website http://www.hgmd.cf.ac.uk/ under the HGMD commercial license using version Q1 of 2021. Due to HGMD commercial licensing, we are not allowed to share these variants publicly, but we provide the following SQL command for reproducibility if you obtain an HGMD commercial license. Variants were downloaded using the following MySQL command from the MySQL import of HGMD 2021q1 SQL dump:

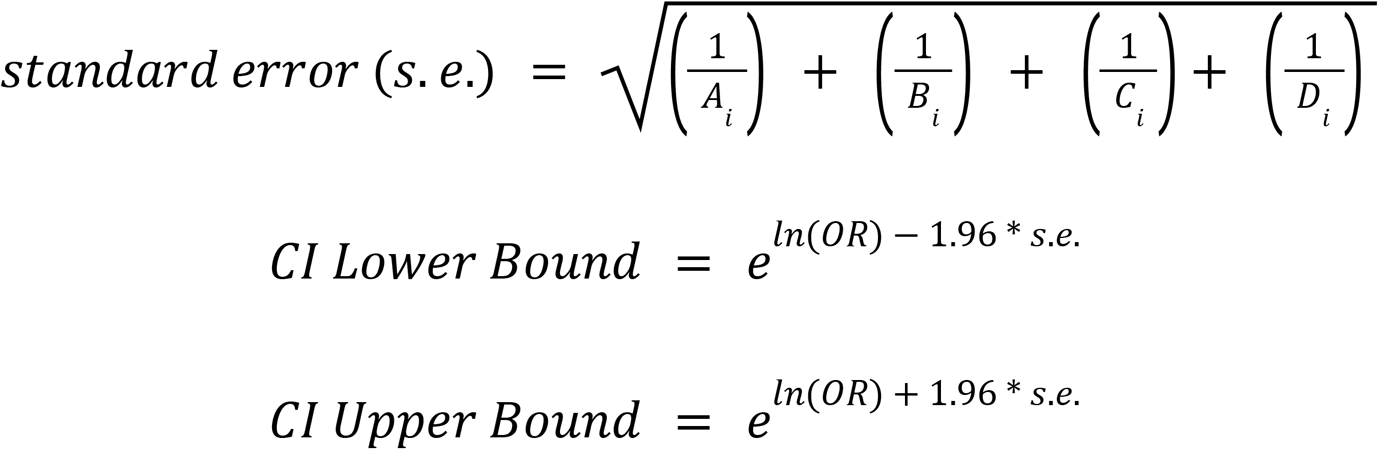

The set of benign variants used in this manuscript are available in **Supplemental Tables S5** of this manuscript.

Scripts, data requirements, and other information that can be used to reproduce the results in this manuscript can be found at the ConSplice manuscript GitHub repo https://github.com/mikecormier/ConSplice-manuscript.

## Acknowledgments

The authors wish to acknowledge Thomas Nicholas, Joe Brown, and Harriet Dashnow for reviews of and feedback on the manuscript.

## Description of Supplemental Data

“Supplemental Material 1” contains Supplemental Figures S1-S19, as well as Supplemental Table S1-S3.

“Supplemental Material 2” contains the manually curated set of pathogenic variants as Supplemental Table S4.

“Supplemental Material 3” contains the set of benign variants used for training and testing the ConSpliceML model as Supplemental Table S5.

