## Supplemental Figures S1-S19 for "Combining genetic constraint with predictions of alternative splicing to prioritize deleterious splicing in rare disease studies"

### Supplemental Material

Supplemental material for the splicing constraint manuscript.

#### Outline


**Supplemental Figures** **2**

Supplemental Figure S1: Splicing constraint model context. 3

Supplemental Figure S2: Patterns of splicing constraint for autosomal genes. 4

Supplemental Figure S3: Patterns of splicing constraint for genes on the autosome and X chromosome. 5

Supplemental Figure S4: Patterns of genic constraint by constraint method. 7

Supplemental Figure S5: Gene expression variation across splicing constraint scores. 9

Supplemental Figure S6: Genic splicing outliers. 11

Supplemental Figure S7: Informative splicing nucleotides by range/window size. 12

Supplemental Figure S8: Spicing constraint around exons. 14

Supplemental Figure S9: Regional constraint around exon features. 15

Supplemental Figure S10: Regional splicing constraint performance by window size. 17

Supplemental Figure S11: SCN1A splicing constraint score distribution. 18

Supplemental Figure S12: Spicing constraint at SCN1A exons. 19

Supplemental Figure S13: Splicing constraint score profiles for deleterious splicing variants identified in RNA-seq data. 20

Supplemental Figure S14: PR curves for ConSpliceML 5-fold cross-validation. 23

Supplemental Figure S15: Splicing prediction and interpretation using the full test set. 24

Supplemental Figure S16: Odds ratio enrichment of pathogenic to benign variants. 25

Supplemental Figure S17: Regional splicing constraint performance for all HGMD pathogenic and benign alternative splicing GTEx variants by model weight. 26

Supplemental Figure S18: Regional splicing constraint performance for non-canonical splice site HGMD pathogenic and benign alternative splicing GTEx variants by model weight. 27

Supplemental Figure S19: Regional splicing constraint performance for non-splice region HGMD pathogenic and benign alternative splicing GTEx variants by model weight. 28

**Supplemental Tables** **29**

Supplemental Table S1: Autosomal splicing substitution rate by reference allele and SpliceAI score range. 29

Supplemental Table S2: X chromosome splicing substitution rate by reference allele and SpliceAI score range 30

Supplemental Table S3: Three SCN1A poison exons and three SCN1A annotated exons 31

**Supplemental Tables as separate files** **32**

Supplemental Table S4: Manually curated set of pathogenic variants 32

Supplemental Table S5: Set of benign variants 32

**Supplemental Figures**

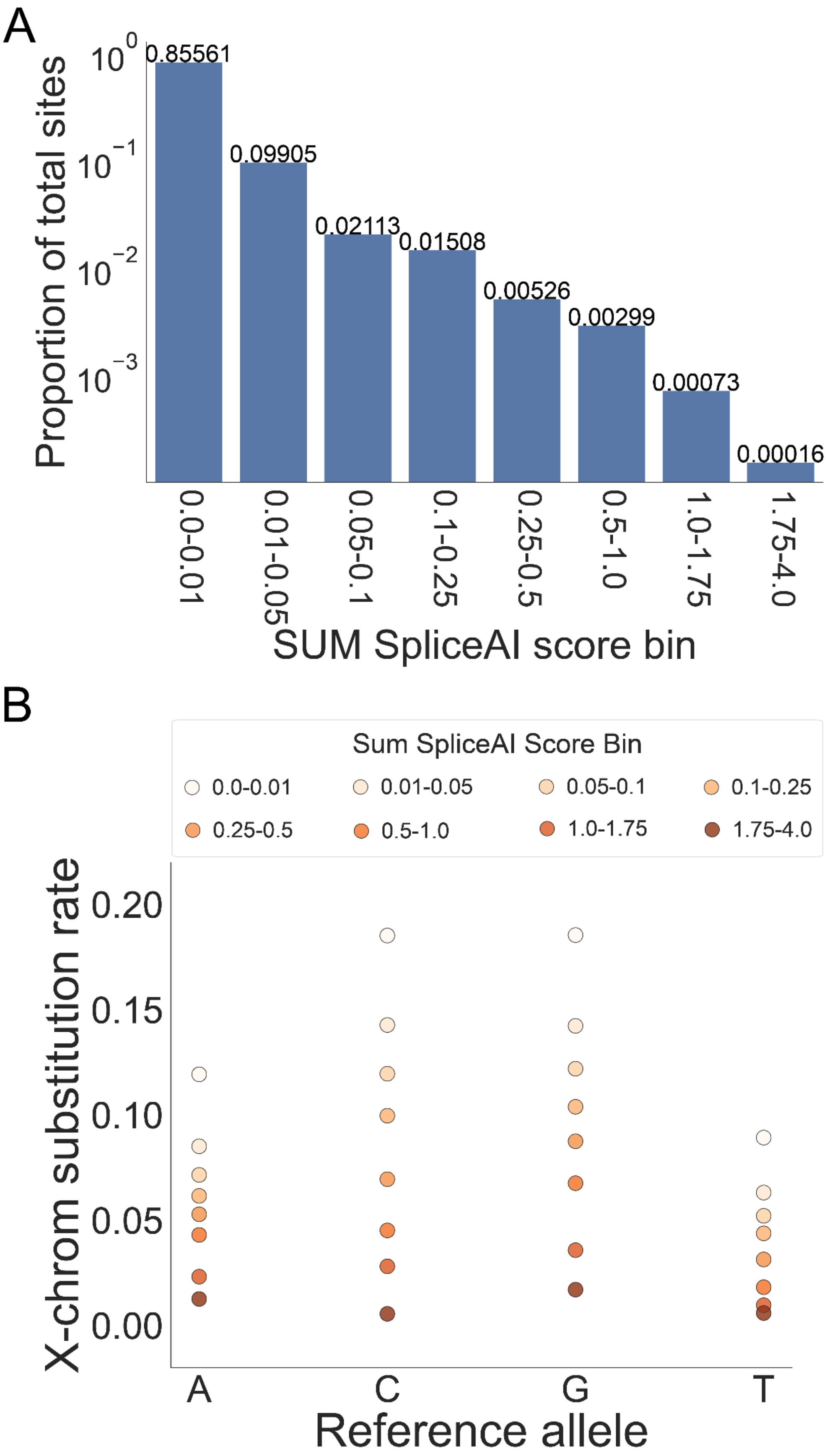

**Supplemental Figure S1: Splicing constraint model context.**

(**A**) The proportion of total sites in protein-coding genes split into the eight SpliceAI score ranges used to create the splicing constraint model. (**B**) The substitution rates for the X chromosome model of splicing constraint. The X chromosome model was created separately from the autosome model due to the difference between the autosome and X chromosome.

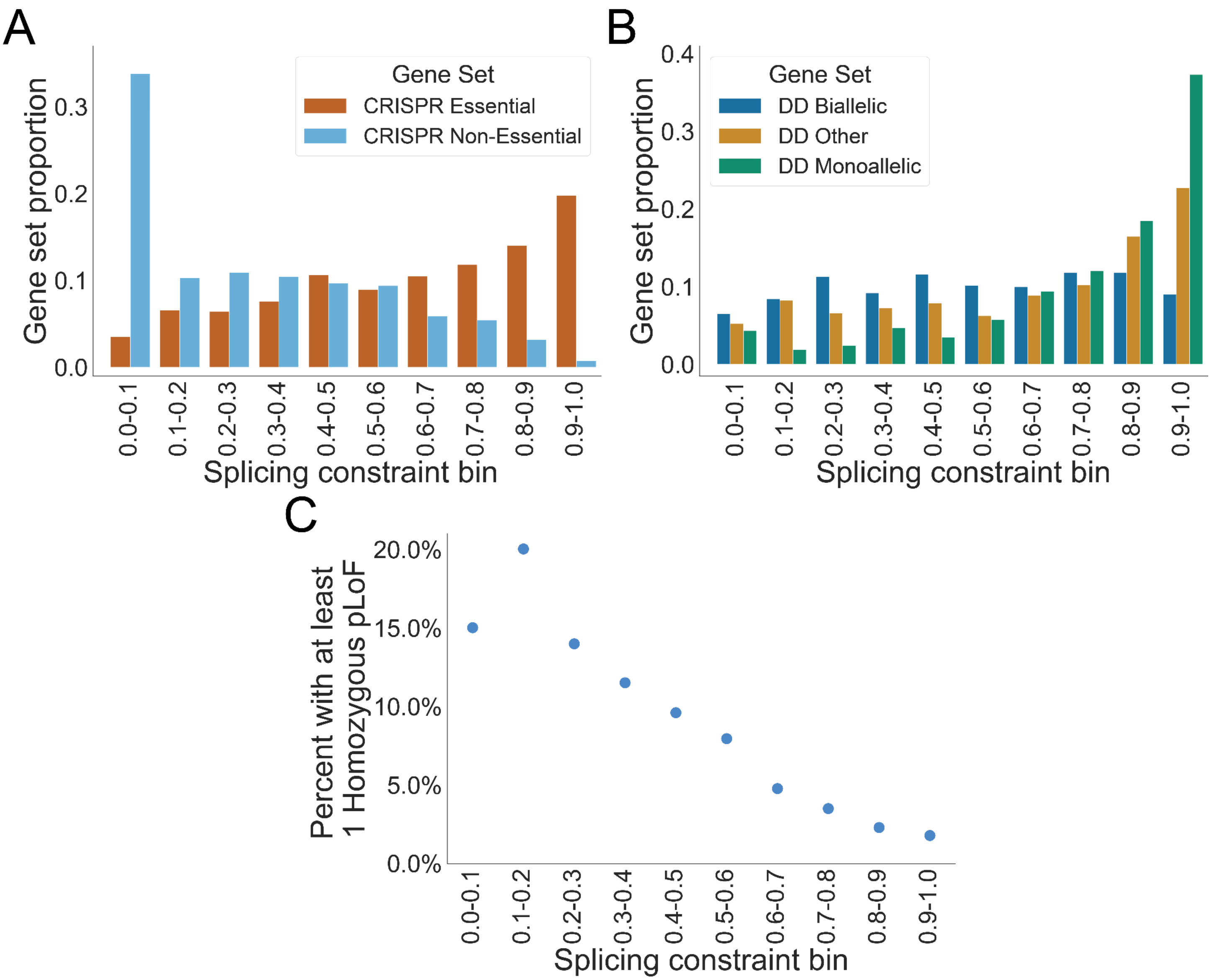


**Supplemental Figure S2: Patterns of splicing constraint for autosomal genes.**

(**A**) The proportion of Essential genes and Non-essential genes across the splicing constraint score range split into deciles. Essential and Non-essential genes were determined by CRISPR screens. (**B**) The proportion of DD/ID biallelic, monoallelic, or other genes split into splicing constraint deciles. DD/ID = developmental delay and intellectual disability. (**C**) The total percent of genes in each splicing constraint decile that are tolerant to at least one homozygous putative loss of function mutation in gnomAD. (Homozygous pLoF tolerant).

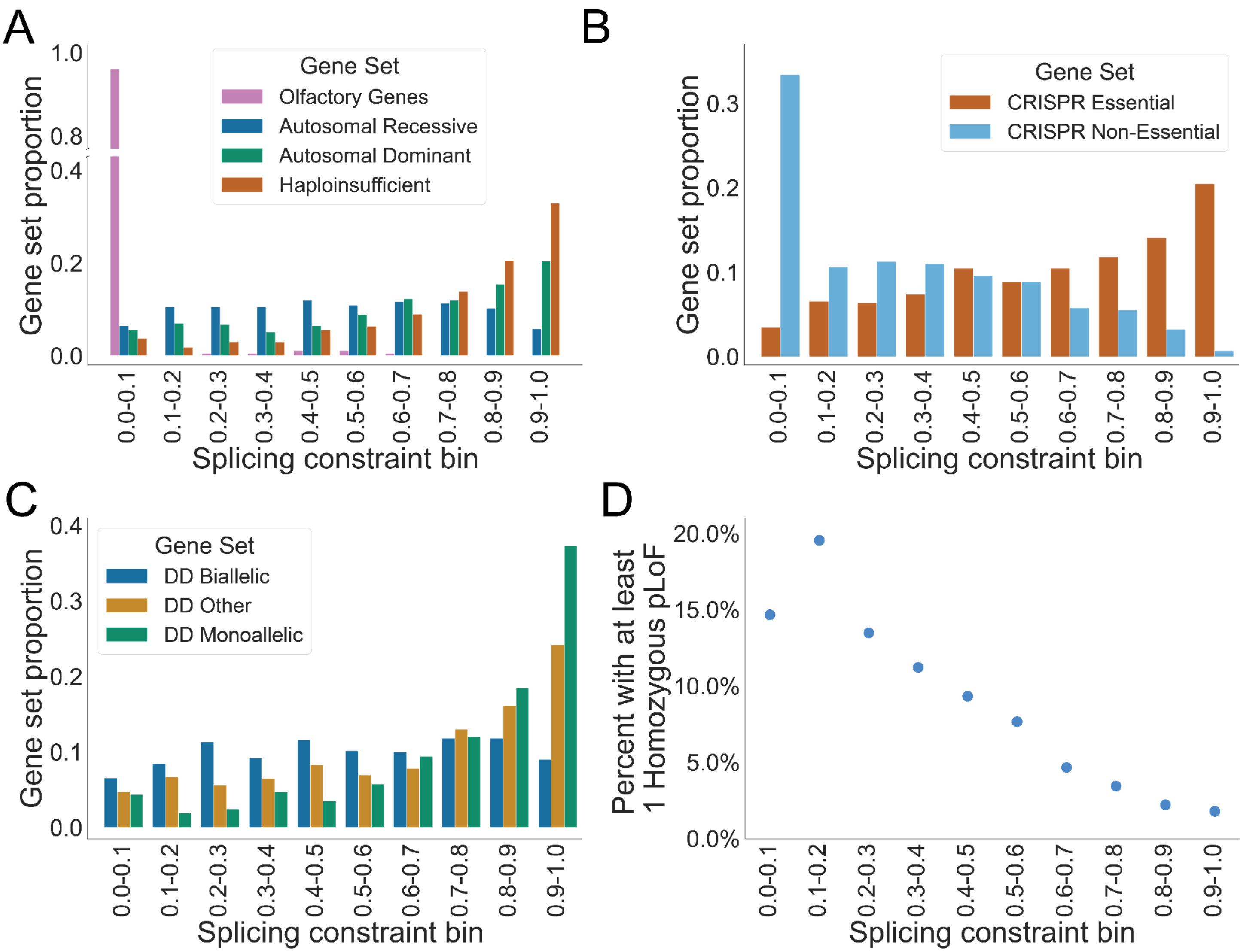


**Supplemental Figure S3: Patterns of splicing constraint for genes on the autosome and X chromosome.**

(**A-D**) Panels A-D are replicated from **Figure 1A** and **Supplemental Figure S2**, with additional genes added from the X chromosome, scored using the X chromosome model of splicing constraint. Therefore, the genes represented in these panels include the autosomal genes scored by the autosomal splicing constraint model and the X chromosome genes scored by the X chromosome splicing constraint model.

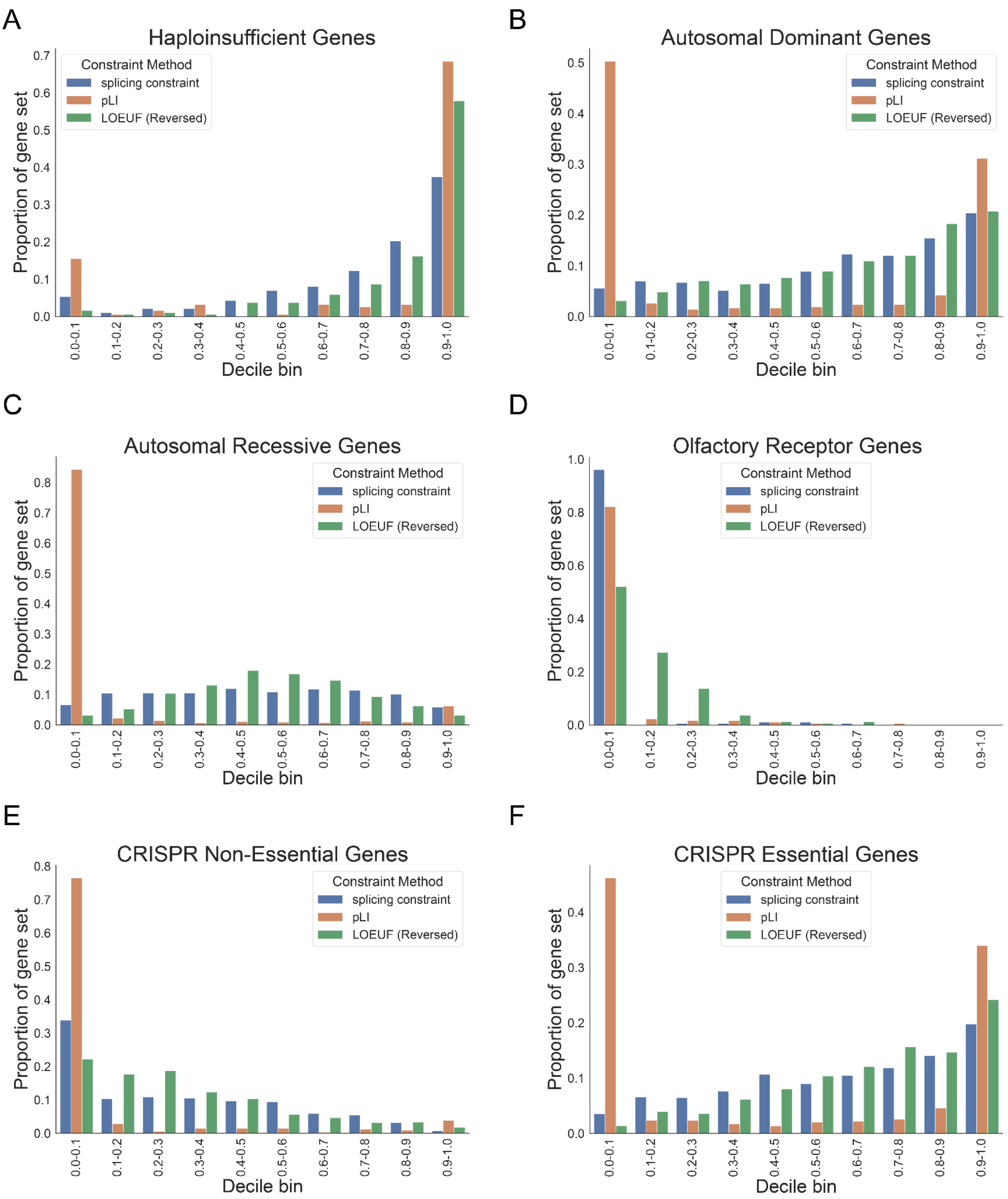
A
Haploinsufficient Genes
B
Autosomal Dominant Genes
C
Autosomal Recessive Genes
D
Olfactory Receptor Genes
E
CRISPR Non-Essential Genes
F
CRISPR Essential Genes
Constraint Method
splicing constraint
pLI
LOEUF (Reversed)
Proportion of gene set
Decile bin
0.0-0.1
0.1-0.2
0.2-0.3
0.3-0.4
0.4-0.5
0.5-0.6
0.6-0.7
0.7-0.8
0.8-0.9
0.9-1.0

**Supplemental Figure S4: Patterns of genic constraint by constraint method.**

The comparison of genic constraint patterns between the splicing constraint metric, pLI, and LEOUF for various gene sets. Genic constraint pattern for (**A**) Haploinsuffiicnet genes, (**B**) Autosomal Dominant genes, (**C**) Autosomal Recessive genes, (**D**) Olfactory Receptor Genes, (**E**) CRISPR non-essential genes, and (**F**) CRISPR essential genes. The color of the bars in each plot represents the constraint method as highlighted by the color legend in each plot.

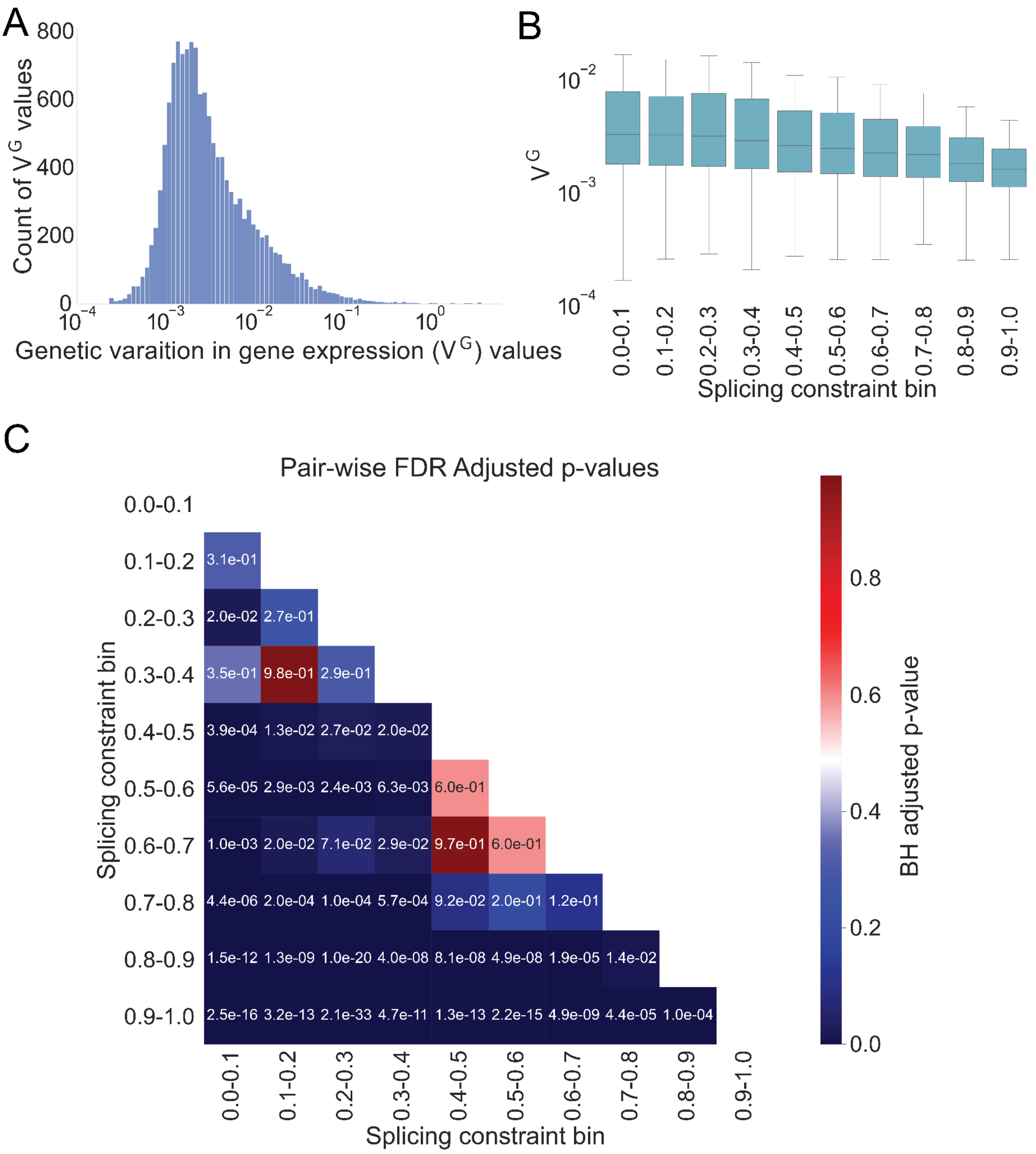
A
Count of V^G values
800
600
400
200
0
10^-4
10^-3
10^-2
10^-1
10^0
Genetic varaition in gene expression (V^G) values
B
V^G
10^-2
10^-3
10^-4
0.0-0.1
0.1-0.2
0.2-0.3
0.3-0.4
0.4-0.5
0.5-0.6
0.6-0.7
0.7-0.8
0.8-0.9
0.9-1.0
Splicing constraint bin
C
Pair-wise FDR Adjusted p-values
0.1-0.2 3.1e-01
0.2-0.3 2.0e-02 2.7e-01
0.3-0.4 3.5e-01 9.8e-01 2.9e-01
0.4-0.5 3.9e-04 1.3e-02 2.7e-02 2.0e-02
0.5-0.6 5.6e-05 2.9e-03 2.4e-03 6.3e-03 6.0e-01
0.6-0.7 1.0e-03 2.0e-02 7.1e-02 2.9e-02 9.7e-01 6.0e-01
0.7-0.8 4.4e-06 2.0e-04 1.0e-04 5.7e-04 9.2e-02 2.0e-01 1.2e-01
0.8-0.9 1.5e-12 1.3e-09 1.0e-20 4.0e-08 8.1e-08 4.9e-08 1.9e-05 1.4e-02
0.9-1.0 2.5e-16 3.2e-13 2.1e-33 4.7e-11 1.3e-13 2.2e-15 4.9e-09 4.4e-05 1.0e-04
Splicing constraint bin
BH adjusted p-value
0.8
0.6
0.4
0.2
0.0

**Supplemental Figure S5: Gene expression variation across splicing constraint scores.**

(**A**) Distribution of $V^G$ values used to compare genic splicing constraint to expression variation. (**B**) $V^G$ distribution by splicing constraint score decile controlling for the number of exons per gene. p-value = **5.99e-26**. Significance determined using ordinary least squares linear regression. Boxes are the same as described in **Figure 2B**. (**C**) The Benjamini-Hochberg FDR adjusted p-values for pair-wise comparison of the $V^G$ distribution in each splicing constraint decile to all other deciles. P-values were determined using a t-test of independence and all p-values were corrected using the Benjamini-Hochberg approach with an alpha value of 0.01. Colors correspond to the adjusted p-value using the heatmap to the right of the p-value matrix. Values in each cell of the matrix correspond to the corrected p-value for the comparison of the splicing constraint decile for that row and column combination.

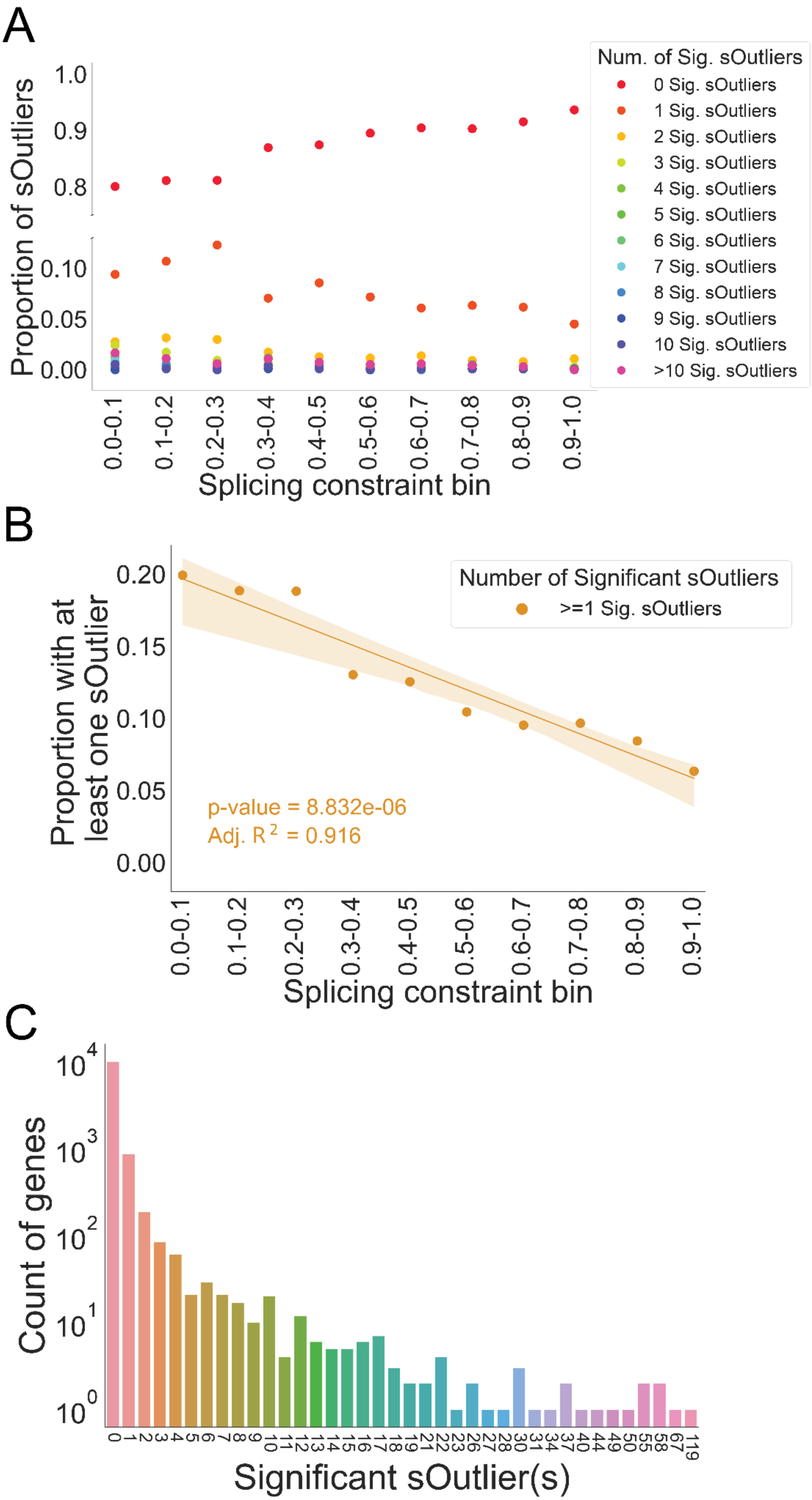
A
Proportion of sOutliers
1.0
0.9
0.8
0.10
0.05
0.00
Num. of Sig. sOutliers
0 Sig. sOutliers
1 Sig. sOutliers
2 Sig. sOutliers
3 Sig. sOutliers
4 Sig. sOutliers
5 Sig. sOutliers
6 Sig. sOutliers
7 Sig. sOutliers
8 Sig. sOutliers
9 Sig. sOutliers
10 Sig. sOutliers
>10 Sig. sOutliers
0.0-0.1
0.1-0.2
0.2-0.3
0.3-0.4
0.4-0.5
0.5-0.6
0.6-0.7
0.7-0.8
0.8-0.9
0.9-1.0
Splicing constraint bin
B
Proportion with at least one sOutlier
0.20
0.15
0.10
0.05
0.00
Number of Significant sOutliers
>=1 Sig. sOutliers
p-value = 8.832e-06
Adj. R 2 = 0.916
0.0-0.1
0.1-0.2
0.2-0.3
0.3-0.4
0.4-0.5
0.5-0.6
0.6-0.7
0.7-0.8
0.8-0.9
0.9-1.0
Splicing constraint bin
C
Count of genes
10^4
10^3
10^2
10^1
10^0
0 1 2 3 4 5 6 7 8 9 10 11 12 13 14 15 16 17 18 19 21 22 23 26 27 28 30 31 34 37 40 44 49 50 55 58 67 119
Significant sOutlier(s)

**Supplemental Figure S6: Genic splicing outliers.**

(**A**) The proportion of genes with splicing outliers in each splicing constraint score decile, split up by the number of significant sOutliers. The combined proportion of sOutliers in a single splicing constraint decile will sum to 1 (100%). (**B**) The proportion of genes in each splicing constraint decile that had one or more significant sOutliers. p-value = **8.832e-06**, Adjusted $R^2$ = **0.916**. Significance determined using ordinary least squares linear regression. Dark orange line represents the linear regression line. Light orange shade represents the 95% confidence interval. (**C**) The distribution of sOutliers by the number of significant sOutliers per gene.

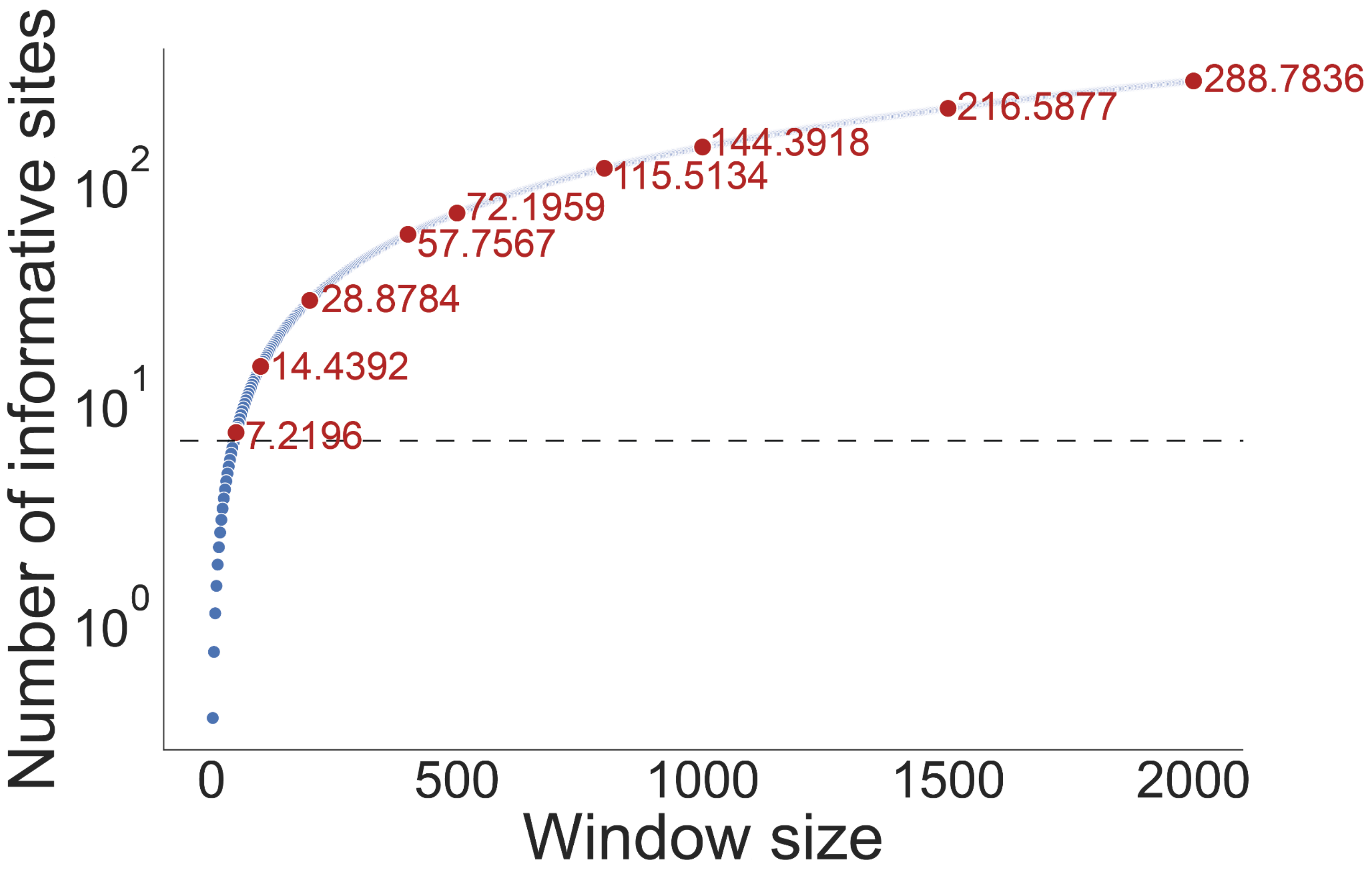


**Supplemental Figure S7: Informative splicing nucleotides by range/window size.**

The number of sites/nucleotides in a given window size that contribute to alternative splicing based on a Poisson distribution using the global proportion of sites with a SpliceAI score above 0.0. Blue dots represent the number of informative sites for a given window size. Red dots represent the window sizes displayed in this manuscript. Red values next to the red dots are the number of informative sites at a given window size. The black horizontal dotted line represents the critical value from the F distribution used to determine a window size that is sufficiently large to capture splicing constraint and an O/E signal (critical value = 6.5415, **Methods**). Any window above the dotted line represents a sufficiently large window to calculate splicing constraint, while any window below the dotted line represents an insufficiently large window. Window sizes corresponding to the red dots from left to right = 50, 100, 200, 400, 500, 800, 1000, 1500, 2000.

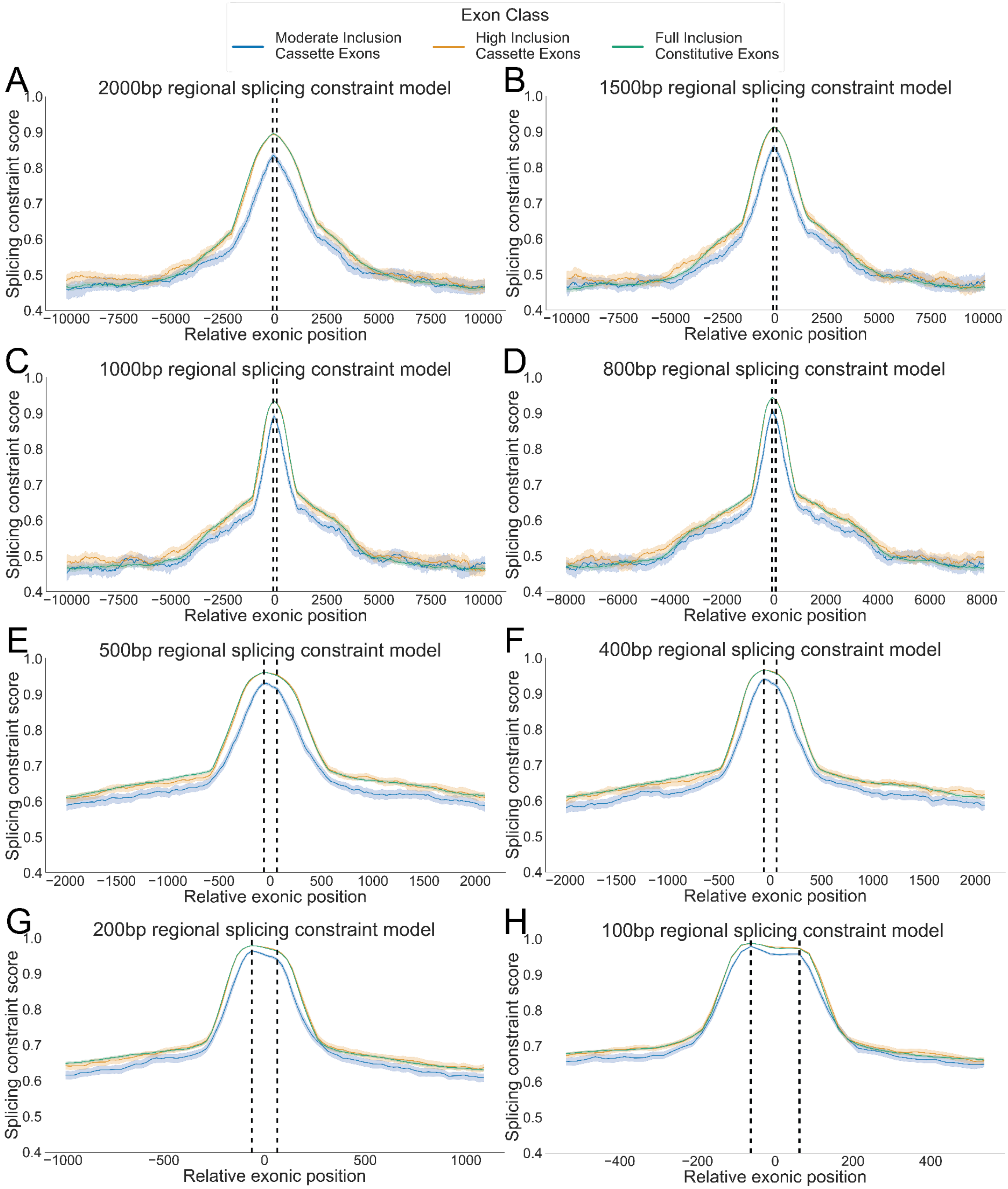
Exon Class
Moderate Inclusion Cassette Exons
High Inclusion Cassette Exons
Full Inclusion Constitutive Exons
A
2000bp regional splicing constraint model
B
1500bp regional splicing constraint model
C
1000bp regional splicing constraint model
D
800bp regional splicing constraint model
E
500bp regional splicing constraint model
F
400bp regional splicing constraint model
G
200bp regional splicing constraint model
H
100bp regional splicing constraint model
Splicing constraint score
Relative exonic position

**Supplemental Figure S8: Spicing constraint around exons.**

The pattern of splicing constraint at and around exons. Exon classes are distinguished by color. Plots are oriented 5’ to 3’ from left to right. Vertical dotted lines represent the start (left) and end (right) positions of exons. (**A**) 2000bp splicing constraint model. (**B**) 1500bp splicing constraint model. (**C**) 1000bp splicing constraint model. (**D**) 800bp splicing constraint model. (**E**) 500bp splicing constraint model. (**F**) 400bp splicing constraint model. (**G**) 200bp splicing constraint model. (**H**) 100bp splicing constraint model. See **Figure 3** for more details.

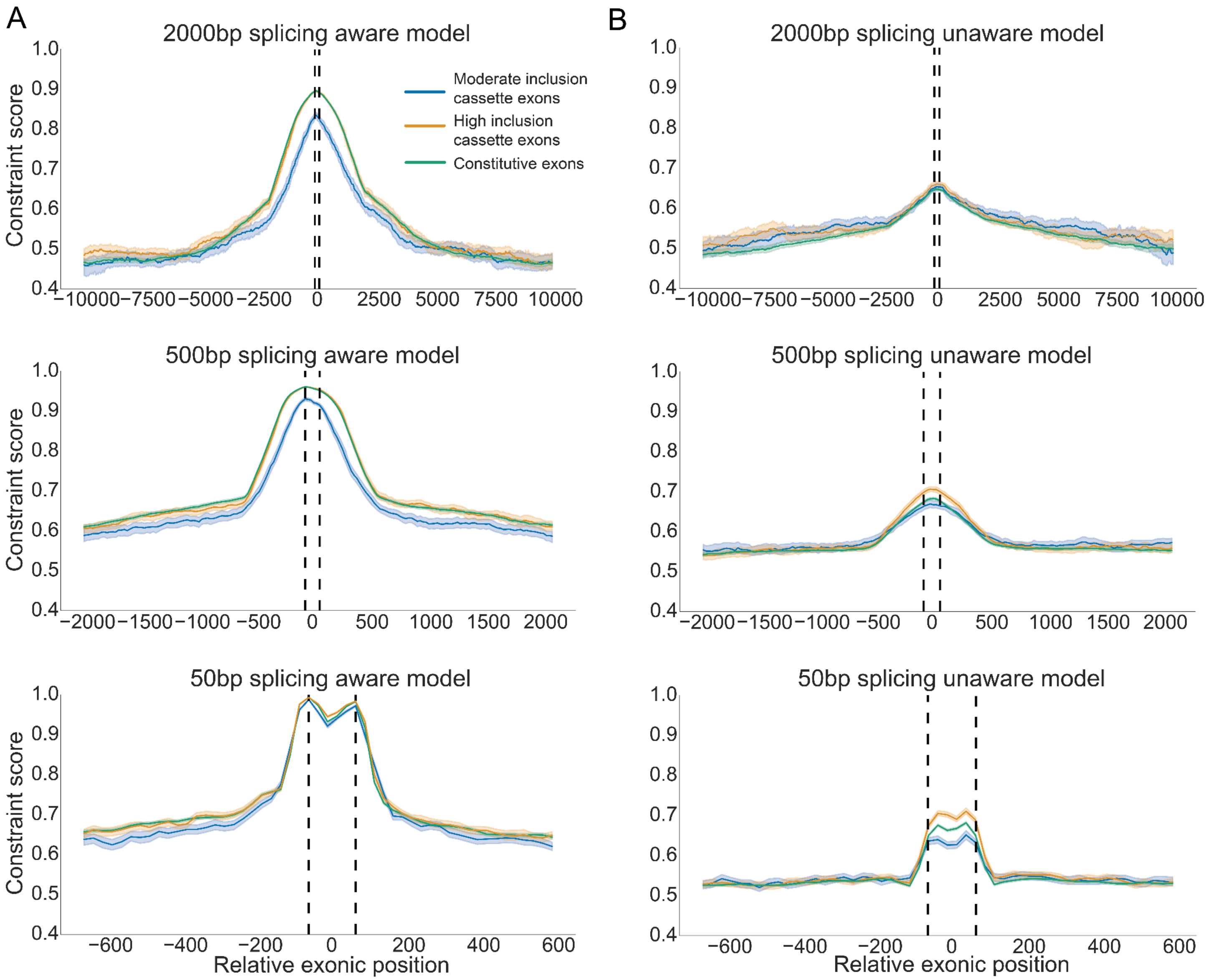


**Supplemental Figure S9: Regional constraint around exon features.**

The regional constraint profiles around exon features for (**A**) the splicing aware model of constraint and (**B**) a generic model of constraint unaware of splicing. See **Figure 3** for more details.

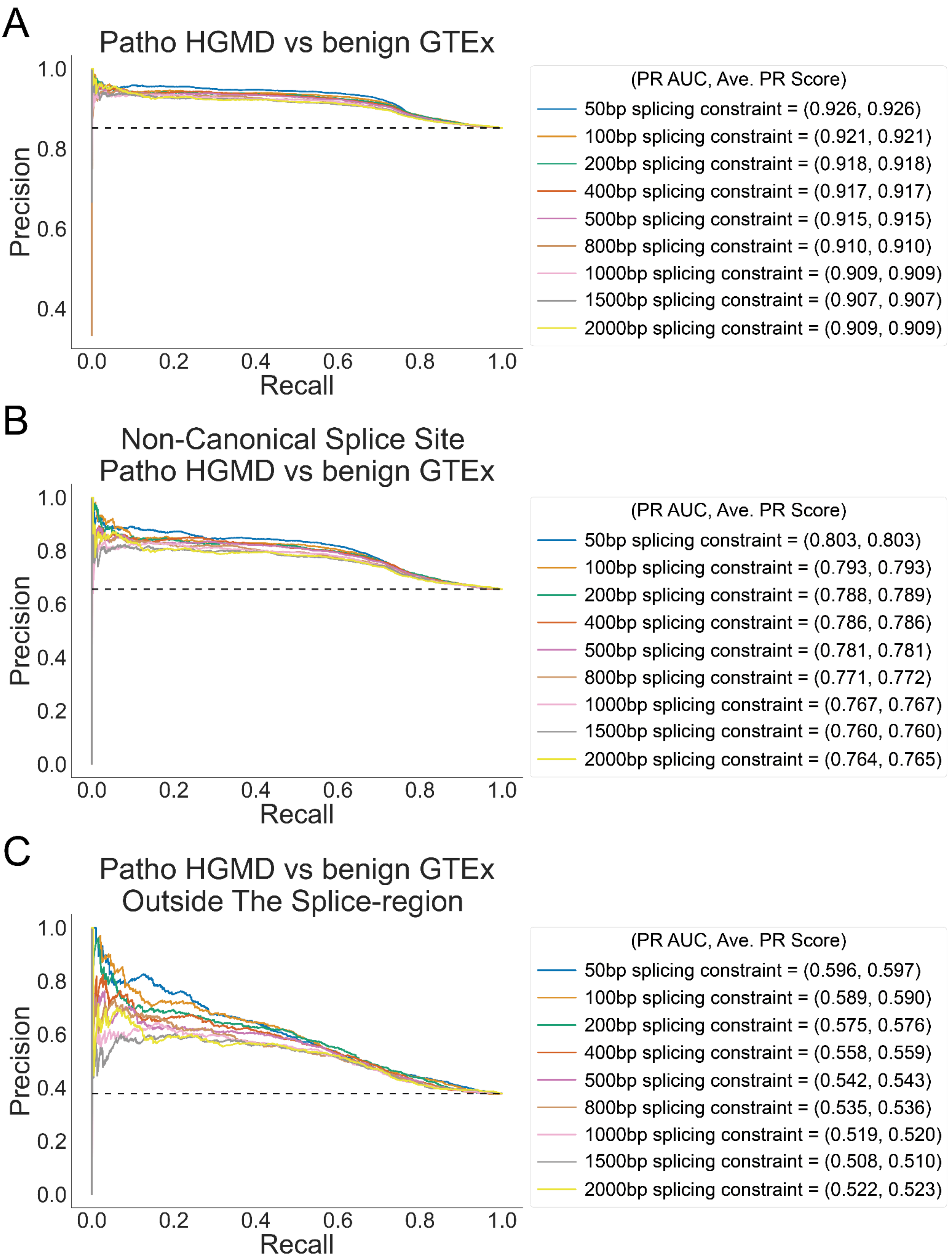
A
Patho HGMD vs benign GTEx
Precision
Recall
(PR AUC, Ave. PR Score)
50bp splicing constraint = (0.926, 0.926)
100bp splicing constraint = (0.921, 0.921)
200bp splicing constraint = (0.918, 0.918)
400bp splicing constraint = (0.917, 0.917)
500bp splicing constraint = (0.915, 0.915)
800bp splicing constraint = (0.910, 0.910)
1000bp splicing constraint = (0.909, 0.909)
1500bp splicing constraint = (0.907, 0.907)
2000bp splicing constraint = (0.909, 0.909)
B
Non-Canonical Splice Site
Patho HGMD vs benign GTEx
Precision
Recall
(PR AUC, Ave. PR Score)
50bp splicing constraint = (0.803, 0.803)
100bp splicing constraint = (0.793, 0.793)
200bp splicing constraint = (0.788, 0.789)
400bp splicing constraint = (0.786, 0.786)
500bp splicing constraint = (0.781, 0.781)
800bp splicing constraint = (0.771, 0.772)
1000bp splicing constraint = (0.767, 0.767)
1500bp splicing constraint = (0.760, 0.760)
2000bp splicing constraint = (0.764, 0.765)
C
Patho HGMD vs benign GTEx
Outside The Splice-region
Precision
Recall
(PR AUC, Ave. PR Score)
50bp splicing constraint = (0.596, 0.597)
100bp splicing constraint = (0.589, 0.590)
200bp splicing constraint = (0.575, 0.576)
400bp splicing constraint = (0.558, 0.559)
500bp splicing constraint = (0.542, 0.543)
800bp splicing constraint = (0.535, 0.536)
1000bp splicing constraint = (0.519, 0.520)
1500bp splicing constraint = (0.508, 0.510)
2000bp splicing constraint = (0.522, 0.523)

**Supplemental Figure S10: Regional splicing constraint performance by window size.**

The performance of the regional splicing constraint model was assessed at different window sizes to empirically identify the best-performing window size to capture focal constraint which can be used to delineate the difference between pathogenic and benign variants. (**A**) Performance of difference splicing constraint models based on region size using the full set of pathogenic variants in the HGMD truth set and all benign validated alternative splicing variants from the GTEx truth set. (**B**) Same as A but excluding variants at the canonical acceptor and donor splice sites. (**C**) Same as A but excluding variants in the splice region. (See **Methods** for more details)

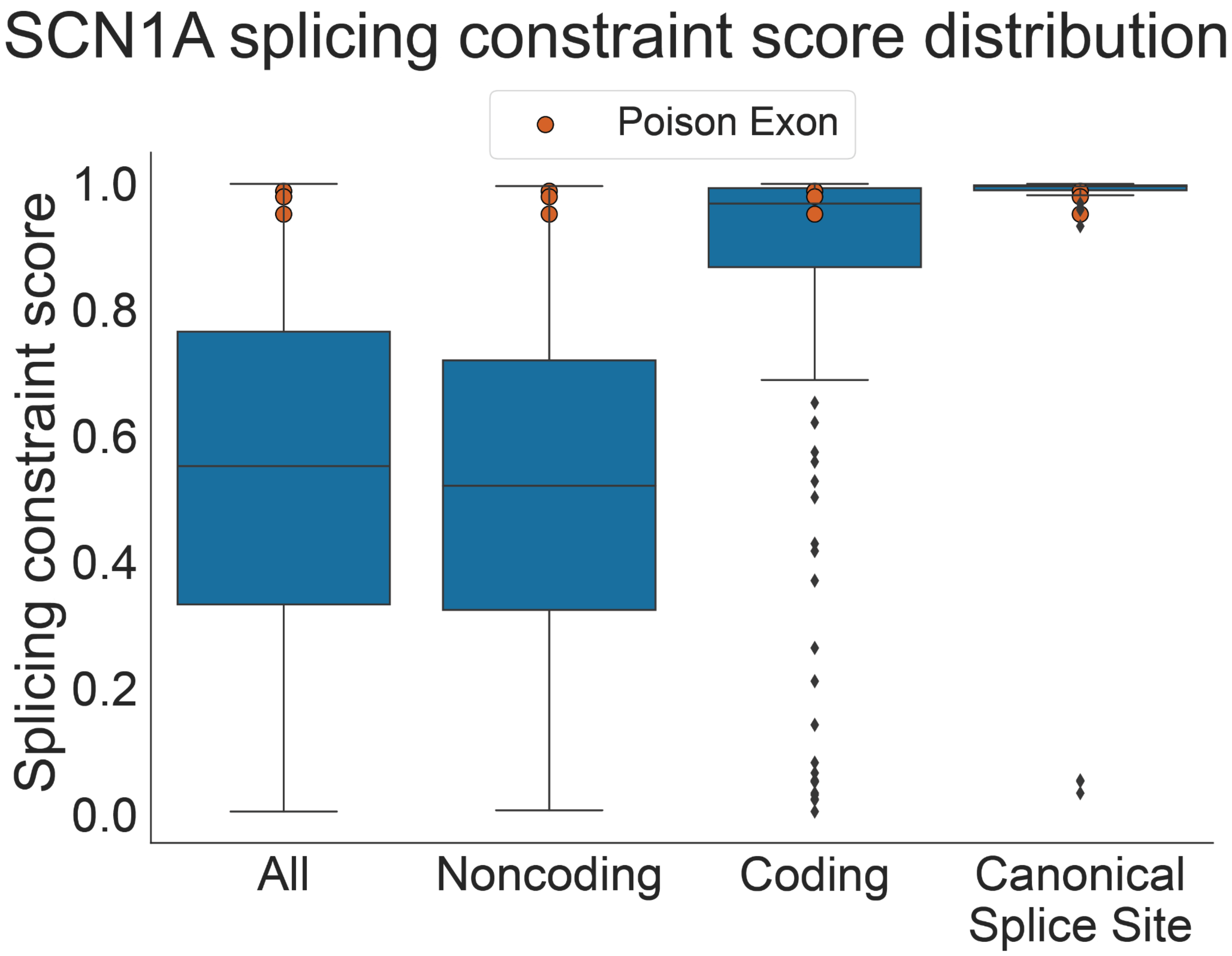


**Supplemental Figure S11: SCN1A splicing constraint score distribution.**

The distribution of splicing constraint scores for All regions in SCN1A, Noncoding regions in SCN1A, Coding regions in SCN1A, and regions that overlap Canonical Splice SItes in SCN1A. The orange dots represent the max splicing constraint score overlapping each of the three poison exons in SCN1A. The splicing constraint around these poison exons resembles that of coding regions and canonical splice sites in SCN1A. The IQR of the boxes ranges from the 25th to 75th percentile. The horizontal black line in each box represents the median splicing constraint value for that distribution. The whiskers are 1.5X the IQR

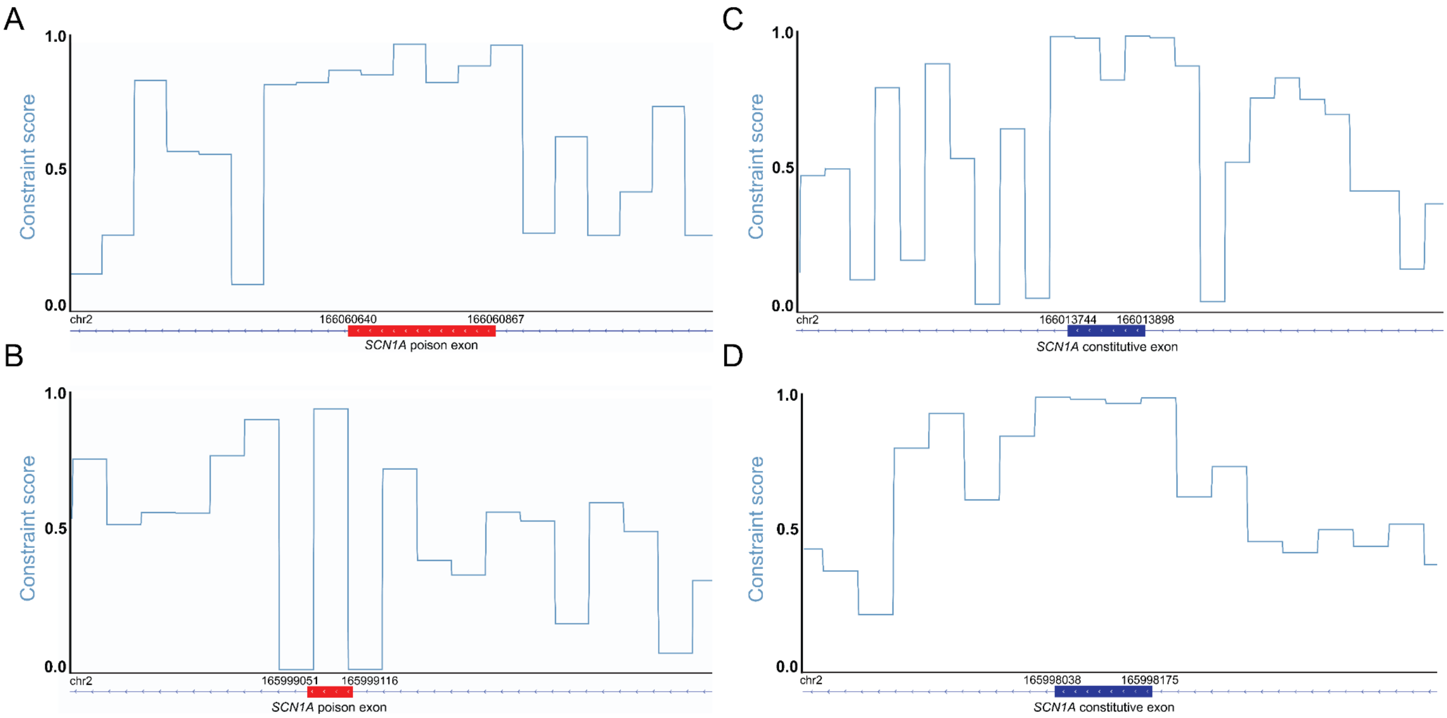


**Supplemental Figure S12: Spicing constraint at SCN1A exons.**

(**A-D**) splicing constraint profile plots of exon features in SCN1A and the corresponding splicing constraint scores. (**A,B**) The splicing constraint profile for two poison exons that cause NMD in SCN1A. Red horizontal bars represent the poison exons in SCN1A. (**C,D**) The splicing constraint profile around two randomly selected canonical exons in SCN1A. Dark blue horizontal lines represent the annotated exons in SCN1A. splicing constraint profiles in A-D are based on the 50bp regional splicing constraint model. The 3' and 5' GRCh38 genomic coordinates for these features are located above the gene track and can be found in **Supplemental Table S3**. For additional details, see **Figure 3**.

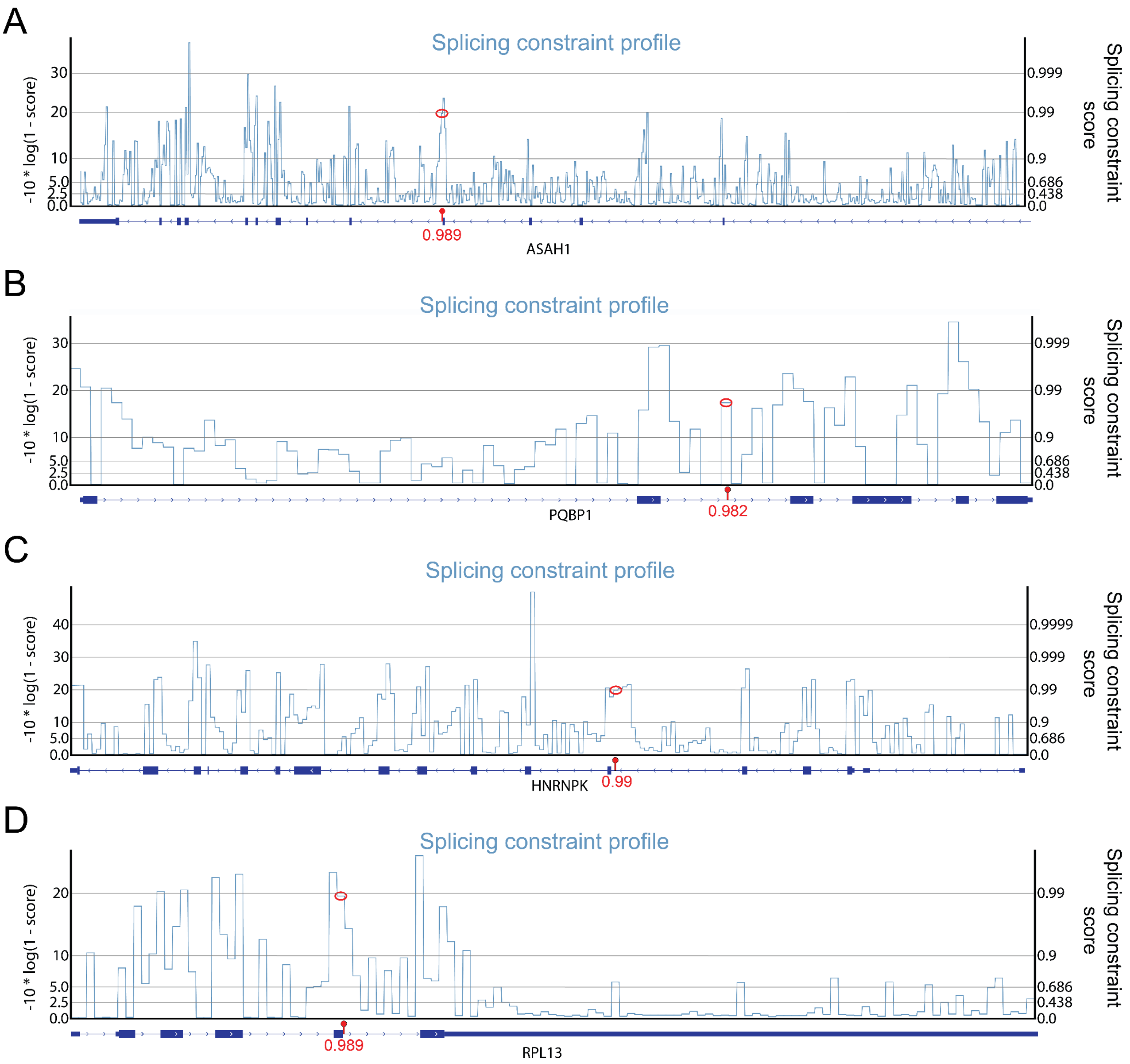


**Supplemental Figure S13: Splicing constraint score profiles for deleterious splicing variants identified in RNA-seq data.**

(**A-D**) Splicing constraint score profile plots displaying pathogenic splicing regions discovered using RNA-seq data, with the pathogenic variant labeled as a red lollipop on the gene track and the corresponding splicing constraint region circled in red on the splicing constraint track. The splicing constraint score is provided below the variant lollipop. The left y-axis shows the log scaled splicing constraint score using -10 * log(1 - splicing constraint score). The right y-axis shows the splicing constraint score for the associated log score. The splicing constraint profile represents the log scaled splicing constraint scores. (**A**) splicing constraint score profile plot for the *ASAH1* gene with a pathogenic splice-altering variant that causes SMAPME. (**B**) Splicing constraint score profile plot for the *PQBP1* gene on the X chromosomes with a pathogenic splice-altering variant that causes RENS1. (**C**) Splicing constraint score profile plot for the *HNRNPK* gene with a pathogenic splice-altering variant that causes AUKS. (**D**) Splicing constraint score profile plot for the *RPL13* gene with a pathogenic splice-altering variant that causes SEMD. SMAPME = Spinal Muscular Atrophy with Progressive Myoclonic Epilepsy. RENS1 = Renpenning Syndrome. AUKS = Au-Kline Syndrome. SEMD = Spondyloepimetaphyseal Dysplasia.

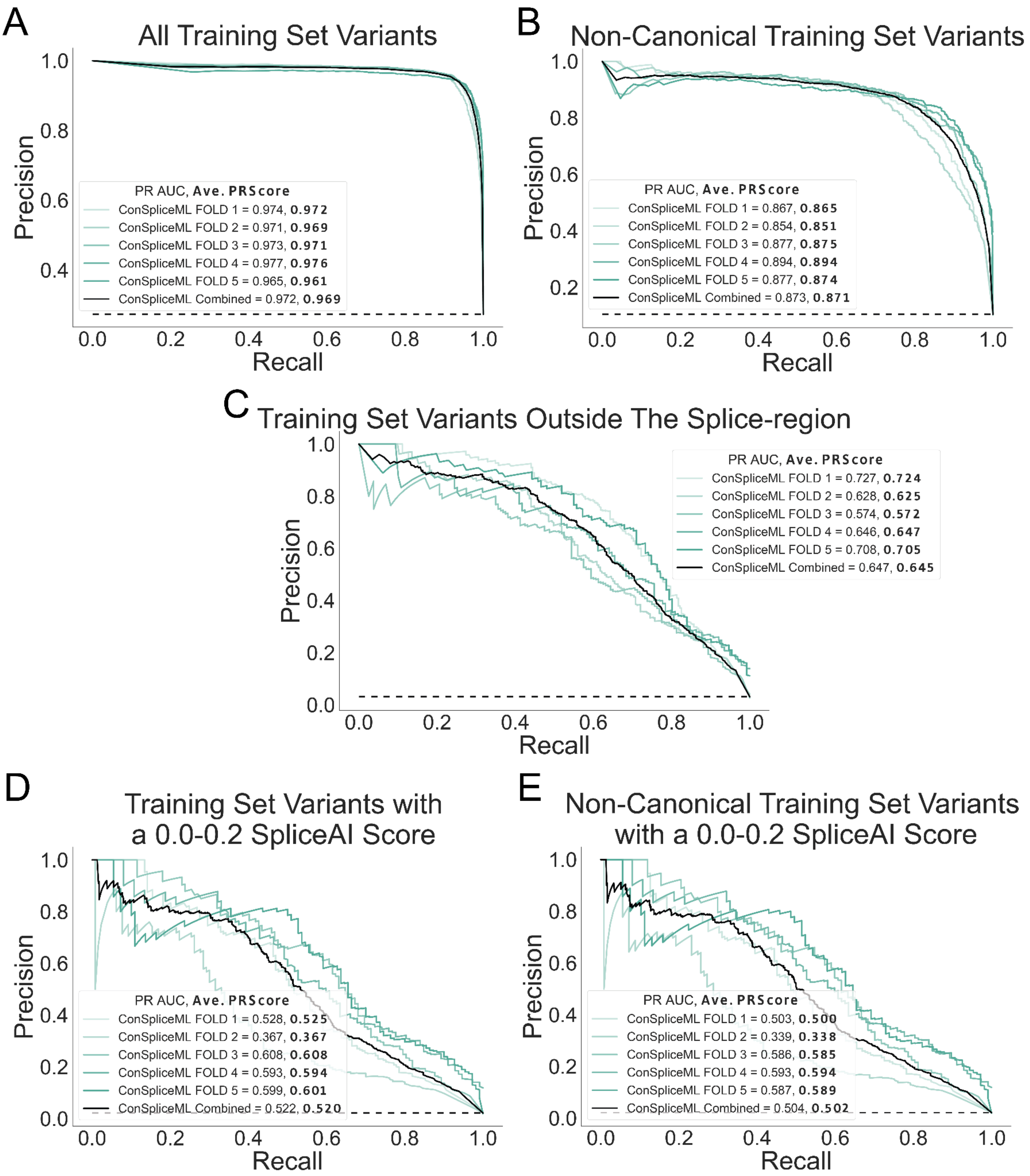
A
All Training Set Variants
Precision
Recall
PR AUC, Ave. PRScore
ConSpliceML FOLD 1 = 0.974, 0.972
ConSpliceML FOLD 2 = 0.971, 0.969
ConSpliceML FOLD 3 = 0.973, 0.971
ConSpliceML FOLD 4 = 0.977, 0.976
ConSpliceML FOLD 5 = 0.965, 0.961
ConSpliceML Combined = 0.972, 0.969
B
Non-Canonical Training Set Variants
Precision
Recall
PR AUC, Ave. PRScore
ConSpliceML FOLD 1 = 0.867, 0.865
ConSpliceML FOLD 2 = 0.854, 0.851
ConSpliceML FOLD 3 = 0.877, 0.875
ConSpliceML FOLD 4 = 0.894, 0.894
ConSpliceML FOLD 5 = 0.877, 0.874
ConSpliceML Combined = 0.873, 0.871
C
Training Set Variants Outside The Splice-region
Precision
Recall
PR AUC, Ave. PRScore
ConSpliceML FOLD 1 = 0.727, 0.724
ConSpliceML FOLD 2 = 0.628, 0.625
ConSpliceML FOLD 3 = 0.574, 0.572
ConSpliceML FOLD 4 = 0.646, 0.647
ConSpliceML FOLD 5 = 0.708, 0.705
ConSpliceML Combined = 0.647, 0.645
D
Training Set Variants with a 0.0-0.2 SpliceAI Score
Precision
Recall
PR AUC, Ave. PRScore
ConSpliceML FOLD 1 = 0.528, 0.525
ConSpliceML FOLD 2 = 0.367, 0.367
ConSpliceML FOLD 3 = 0.608, 0.608
ConSpliceML FOLD 4 = 0.593, 0.594
ConSpliceML FOLD 5 = 0.599, 0.601
ConSpliceML Combined = 0.522, 0.520
E
Non-Canonical Training Set Variants with a 0.0-0.2 SpliceAI Score
Precision
Recall
PR AUC, Ave. PRScore
ConSpliceML FOLD 1 = 0.503, 0.500
ConSpliceML FOLD 2 = 0.339, 0.338
ConSpliceML FOLD 3 = 0.586, 0.585
ConSpliceML FOLD 4 = 0.593, 0.594
ConSpliceML FOLD 5 = 0.587, 0.589
ConSpliceML Combined = 0.504, 0.502

**Supplemental Figure S14: PR curves for ConSpliceML 5-fold cross-validation.**

(**A-E**) The Precision-Recall curves for each of the five cross-validation folds used to train ConSpliceML. The performance of each fold is designated by a shade of gray. The black curve represents the combined/averaged performance of ConSpliceML across all five folds. (**A**) All variants in the training set. (**B**) All but canonical splice site variants in the training set. (**C**) All variants in the training set outside of the splice region. (**D**) All variants in the filtered training set with a SpliceAI score between 0.0 and 0.2. (E) Non-canonical variants in the training set with a SpliceAI score between 0.0 and 0.2.

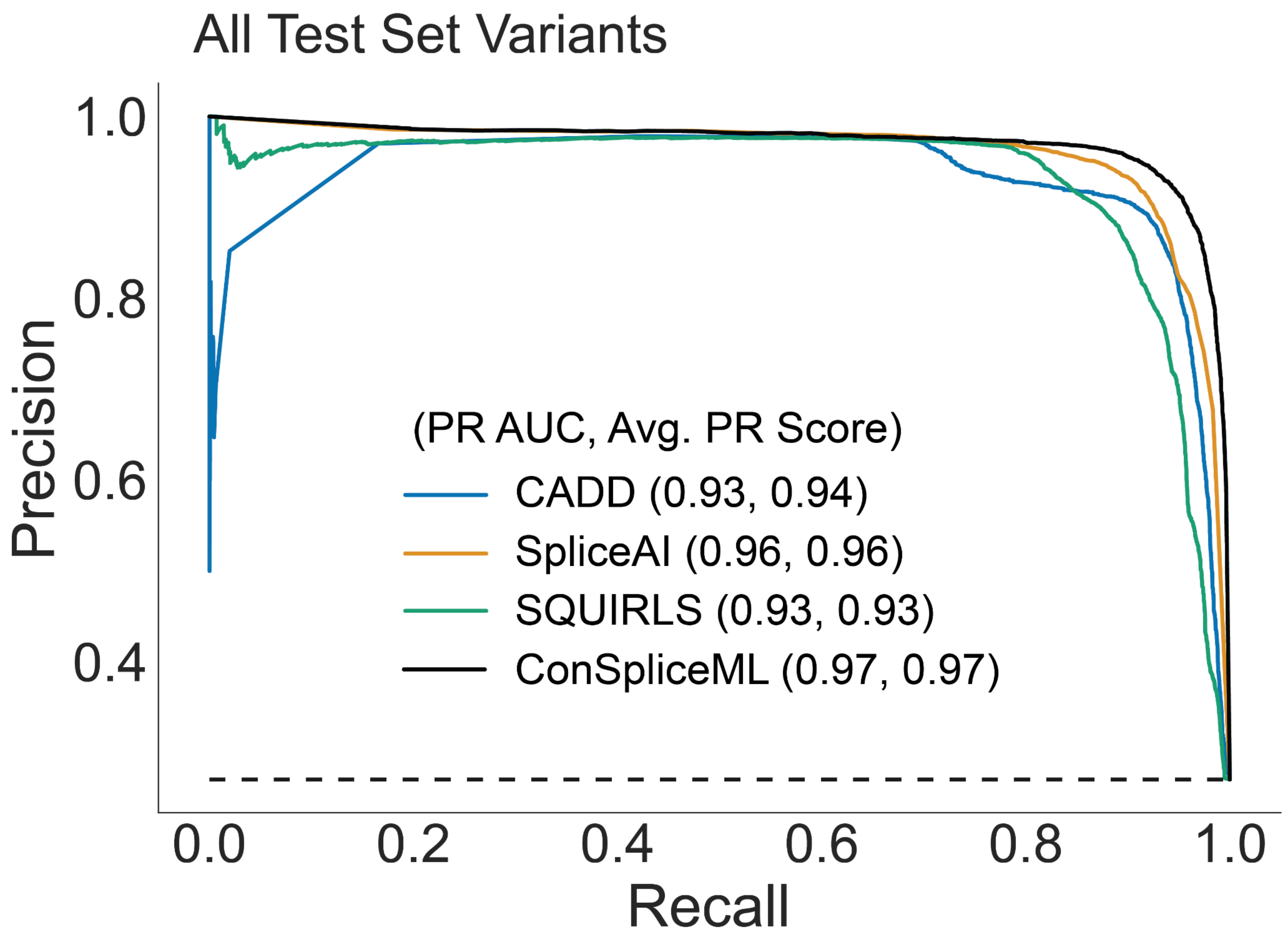


**Supplemental Figure S15: Splicing prediction and interpretation using the full test set.**

Precision-Recall curve depicting the performance of CADD, SpliceAI, SQUIRLS, and ConSpliceML in differentiating pathogenic splicing variants from benign variants using all variants in the test set. 7,269 pathogenic variants. 19,592 benign variants.

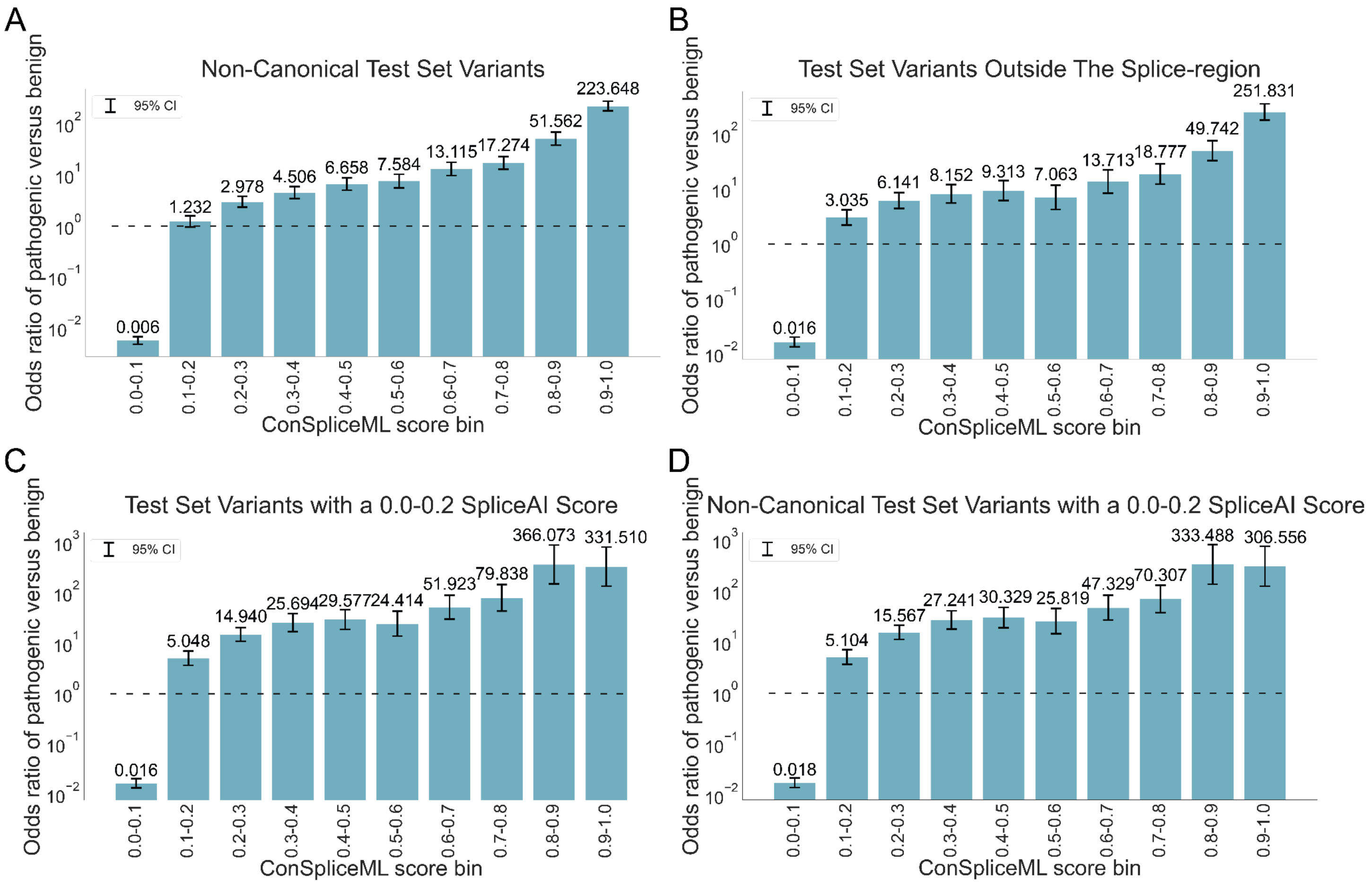


**Supplemental Figure S16: Odds ratio enrichment of pathogenic to benign variants.**

(**A-D**) The odds ratio enrichment of pathogenic to benign variants across the ConSpliceML deciles. Odds ratio values are highlighted in black above each bar. Error bars represent the 95% CI. The dotted horizontal line represents an odds ratio at 1. Variants used to calculate the odds ratios are from the test set. (**A**) Odds ratio enrichment for all non-canonical variants in the test set. (**B**) Odds ratio enrichment for all variants in the test set outside of the splice-region. (**C**) All variants in the test set with a SpliceAI score between 0.0 and 0.2. (**D**) All non-canonical variants in the test set with a SpliceAI score between 0.0 and 0.2.

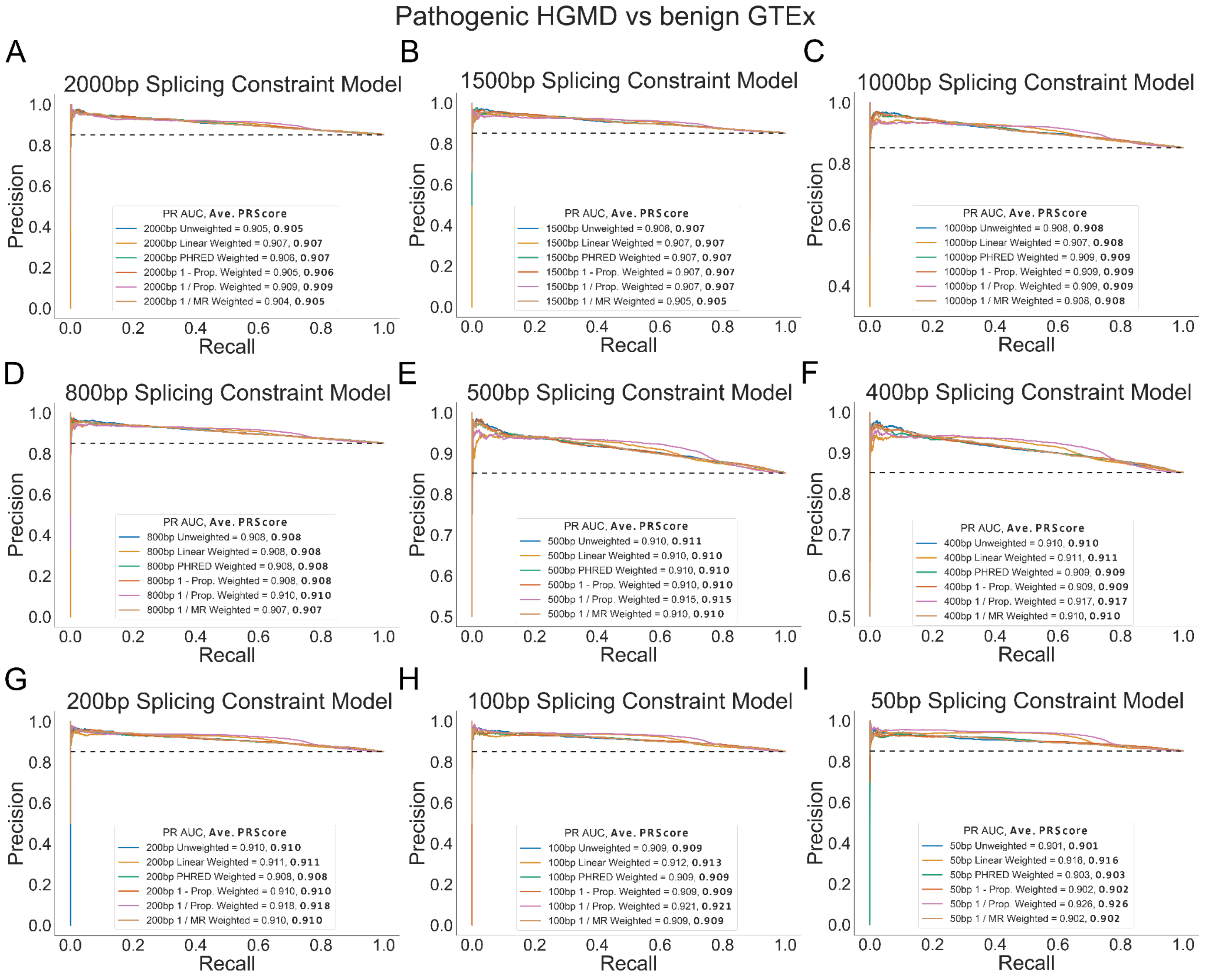


**Supplemental Figure S17: Regional splicing constraint performance for all HGMD pathogenic and benign alternative splicing GTEx variants by model weight.**

The regional splicing constraint performance using the HGMD pathogenic variants and the validated benign splice altering variants from the benign set for each of the six alternative splicing likelihood weights. (**A**) PR curve using the 2000bp splicing constraint model for each of the 6 splicing likelihood weights. (**B**) 1500bp splicing constraint model. (**C**) 1000bp splicing constraint model. (**D**) 800bp splicing constraint model. (**E**) 500bp splicing constraint model. (**F**) 400bp splicing constraint model. (**G**) 200bp splicing constraint model. (**H**) 100bp splicing constraint model. (**I**) 50bp splicing constraint model.

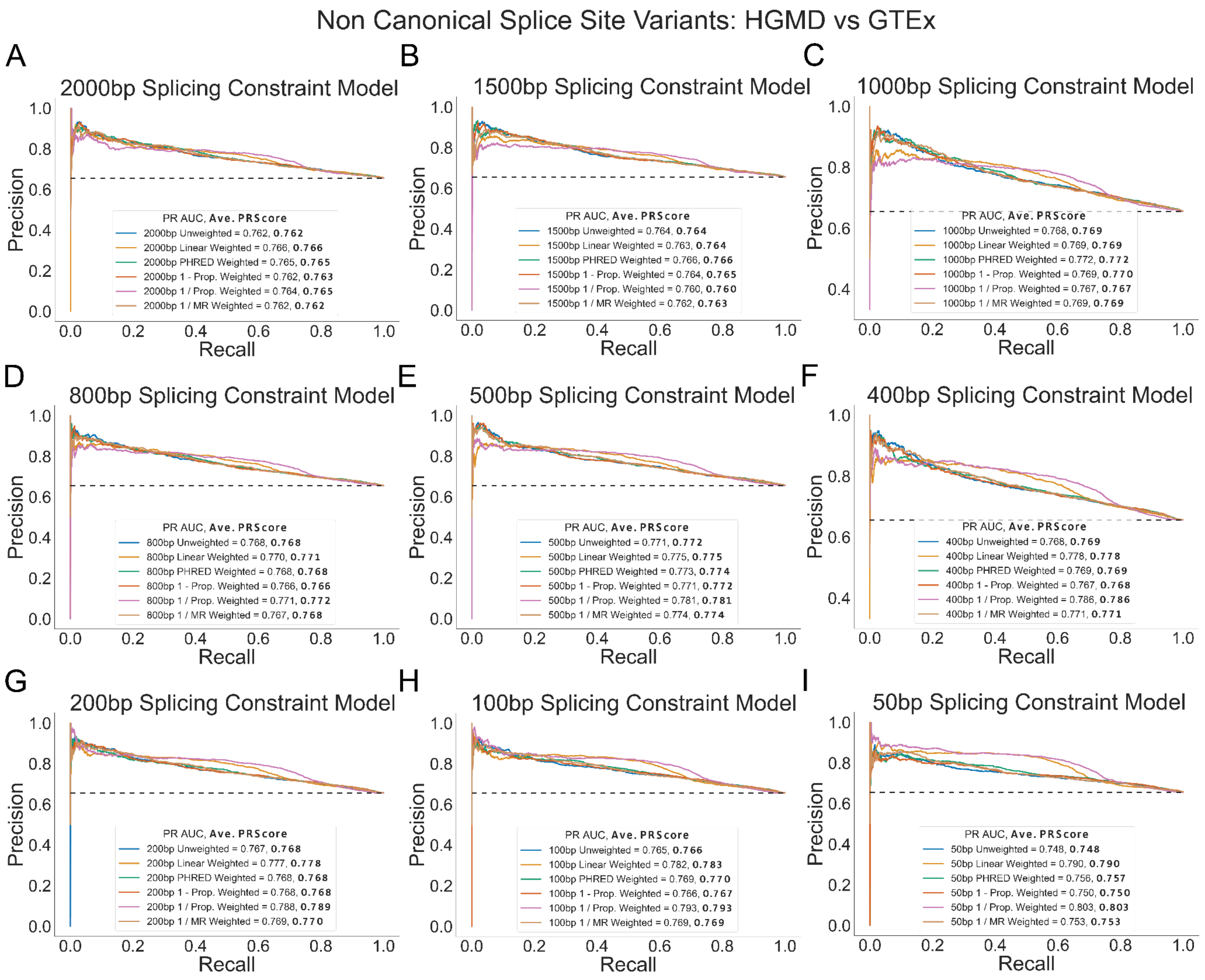


**Supplemental Figure S18: Regional splicing constraint performance for non-canonical splice site HGMD pathogenic and benign alternative splicing GTEx variants by model weight.**

The regional splicing constraint performance for non-canonical splice site variants in the HGMD pathogenic variants and the validated benign splice altering variants from the benign set for each of the six alternative splicing likelihood weights. (**A**) PR curve using the 2000bp splicing constraint model for each of the 6 splicing likelihood weights. (**B**) 1500bp splicing constraint model. (**C**) 1000bp splicing constraint model. (**D**) 800bp splicing constraint model. (**E**) 500bp splicing constraint model. (**F**) 400bp splicing constraint model. (**G**) 200bp splicing constraint model. (**H**) 100bp splicing constraint model. (**I**) 50bp splicing constraint model.

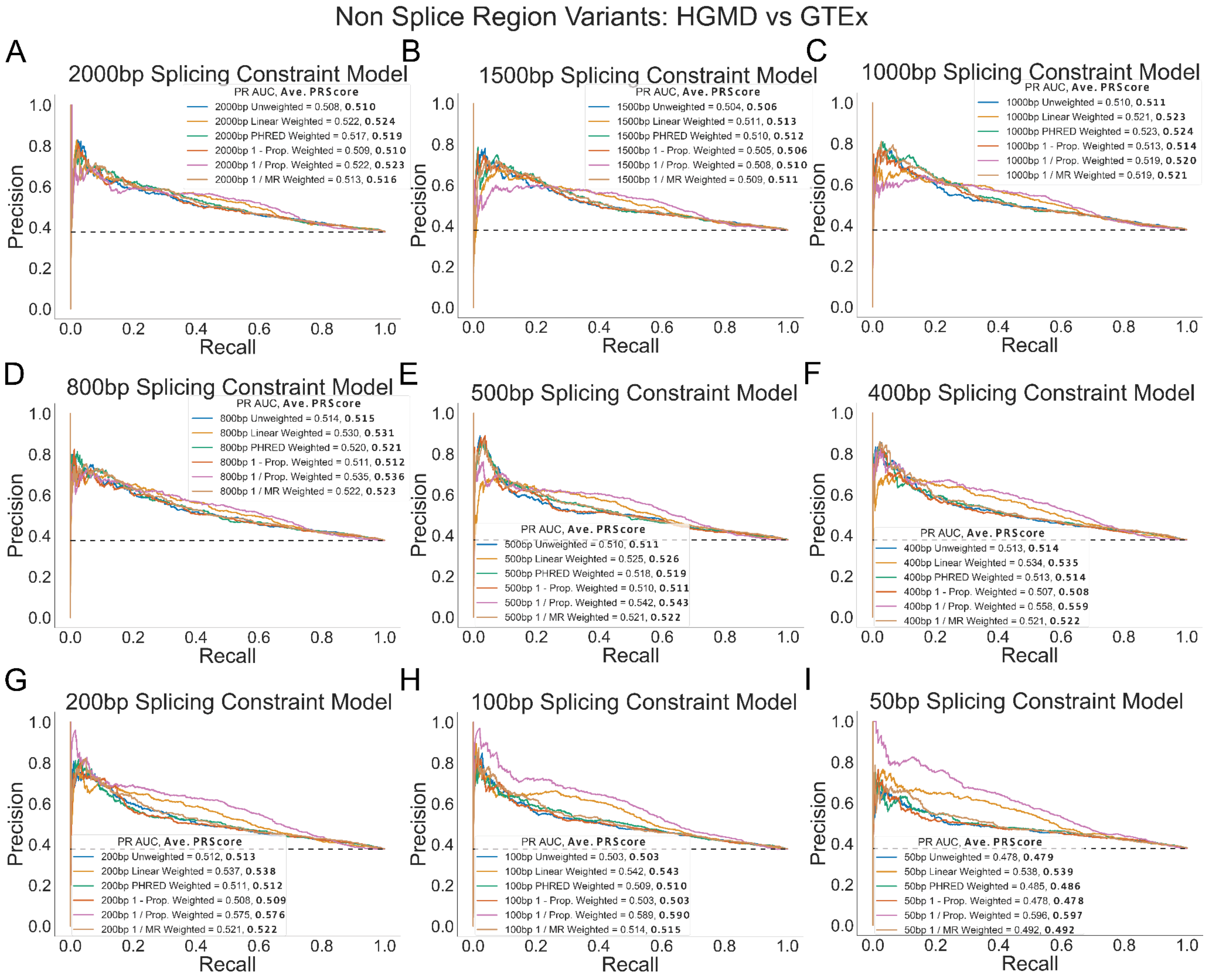


**Supplemental Figure S19: Regional splicing constraint performance for non-splice region HGMD pathogenic and benign alternative splicing GTEx variants by model weight.**

The regional splicing constraint performance for variants outside of the splice region in the HGMD pathogenic variants and the validated benign splice altering variants from the benign set for each of the six alternative splicing likelihood weights. (**A**) PR curve using the 2000bp splicing constraint model for each of the 6 splicing likelihood weights. (**B**) 1500bp splicing constraint model. (**C**) 1000bp splicing constraint model. (**D**) 800bp splicing constraint model. (**E**) 500bp splicing constraint model. (**F**) 400bp splicing constraint model. (**G**) 200bp splicing constraint model. (**H**) 100bp splicing constraint model. (**I**) 50bp splicing constraint model.

**Supplemental Tables**

| | **0.00-0.01** | **0.01-0.05** | **0.05-0.1** | **0.1-0.25** | **0.25-0.5** | **0.5-1.0** | **1.0-1.75** | **1.75-4.0** |
|---|---|---|---|---|---|---|---|---|
| **A** | 0.17859 | 0.13300 | 0.11347 | 0.09880 | 0.08770 | 0.07821 | 0.06363 | 0.04451 |
| **C** | 0.27343 | 0.22104 | 0.18902 | 0.15996 | 0.12382 | 0.08906 | 0.05558 | 0.03576 |
| **G** | 0.26586 | 0.21535 | 0.18851 | 0.16462 | 0.14008 | 0.12574 | 0.09538 | 0.05785 |
| **T** | 0.13592 | 0.10079 | 0.08471 | 0.07232 | 0.05445 | 0.03651 | 0.02357 | 0.01394 |

**Supplemental Table S1: Autosomal splicing substitution rate by reference allele and SpliceAI score range.**

| | **0.00-0.01** | **0.01-0.05** | **0.05-0.1** | **0.1-0.25** | **0.25-0.5** | **0.5-1.0** | **1.0-1.75** | **1.75-4.0** |
|---|---|---|---|---|---|---|---|---|
| **A** | 0.11931 | 0.08512 | 0.07153 | 0.06157 | 0.05284 | 0.04304 | 0.02319 | 0.01261 |
| **C** | 0.18524 | 0.14275 | 0.11962 | 0.09967 | 0.06949 | 0.04517 | 0.02809 | 0.00554 |
| **G** | 0.18548 | 0.14235 | 0.12199 | 0.10389 | 0.08751 | 0.06759 | 0.03579 | 0.01703 |
| **T** | 0.08926 | 0.06315 | 0.05209 | 0.04377 | 0.03136 | 0.01818 | 0.00964 | 0.00591 |

**Supplemental Table S2: X chromosome splicing substitution rate by reference allele and SpliceAI score range**

| chr | genomic start position (GRCh38) | genomic end position (GRCh38) | exon size (in nucleotides) | poison exon (yes/no) | biotype | MAX overlapping splicing constraint score |
|---|---|---|---|---|---|---|
| 2 | 165998038 | 165998175 | 138 | n | Coding | 0.997 |
| 2 | 165999051 | 165999116 | 66 | y | NMD | 0.952 |
| 2 | 166007230 | 166007293 | 64 | y | NMD | 0.988 |
| 2 | 166013744 | 166013898 | 155 | n | Coding | 0.999 |
| 2 | 166051719 | 166051988 | 270 | n | Coding | 0.995 |
| 2 | 166060640 | 166060867 | 228 | y | NMD | 0.979 |

**Supplemental Table S3: Three SCN1A poison exons and three SCN1A annotated exons**

**<u>Supplemental Tables as separate files</u>**

**Supplemental Table S4: Manually curated set of pathogenic variants**

**Supplemental Table S5: Set of benign variants**
